# High Light and High Temperature Reduce Photosynthesis via Different Mechanisms in the C_4_ Model *Setaria viridis*

**DOI:** 10.1101/2021.02.20.431694

**Authors:** Cheyenne M. Anderson, Erin M. Mattoon, Ningning Zhang, Eric Becker, William McHargue, Jiani Yang, Dhruv Patel, Oliver Dautermann, Scott A. M. McAdam, Tonantzin Tarin, Sunita Pathak, Tom J. Avenson, Jeffrey Berry, Maxwell Braud, Krishna K. Niyogi, Margaret Wilson, Dmitri A. Nusinow, Rodrigo Vargas, Kirk J. Czymmek, Andrea L. Eveland, Ru Zhang

## Abstract

C_4_ plants frequently experience damaging high light (HL) and high temperature (HT) conditions in native environments, which reduce growth and yield. However, the mechanisms underlying these stress responses in C_4_ plants have been under-explored, especially the coordination between mesophyll (M) and bundle sheath (BS) cells. We investigated how the C_4_ model plant *Setaria viridis* responded to a four-hour HL or HT treatment at the photosynthetic, transcriptomic, and ultrastructural levels. Although we observed a comparable reduction of photosynthetic efficiency in HL- or HT-treated leaves, detailed analysis of multi-level responses revealed important differences in key pathways and M/BS specificity responding to HL and HT. We provide a systematic analysis of HL and HT responses in *S. viridis*, reveal different acclimation strategies to these two stresses in C_4_ plants, discover unique light/temperature responses in C_4_ plants in comparison to C_3_ plants, and identify potential targets to improve abiotic stress tolerance in C_4_ crops.

## Introduction

Several of the world’s most economically important staple crops utilize C_4_ photosynthesis, including *Zea mays* and *Sorghum bicolor*. C_4_ photosynthesis concentrates CO_2_ around Rubisco (ribulose-1,5-bisphosphate carboxylase/oxygenase) by employing biochemical reactions within mesophyll (M) and bundle sheath (BS) cells^1,2^. The high local concentration of CO_2_ near Rubisco favors carbon fixation over photorespiration, which is initiated by the oxygenase activity of Rubisco^1,3^. C_4_ photosynthesis is hypothesized to have been selected by low CO_2_, high light (HL), and high temperature (HT) conditions^4,5^. C_4_ plants typically exhibit higher photosynthetic and water-use efficiencies than their C_3_ counterparts under HL or HT^6^. However, C_4_ crops experience more frequent damaging HL or HT stresses in their natural environments than C_3_ crops, with reduced C_4_ crop yield regularly occurring in warmer regions^7^. As mean global temperatures continue to increase, maize yields are estimated to decrease between 4 and 12% for each temperature increase in degree Celsius^7^. Photosynthesis in maize leaves is inhibited at leaf temperature above 38°C. Recent data from 408 sorghum cultivars shows that breeding efforts over the last few decades have developed high-yielding sorghum cultivars with considerable variability in heat resilience and even the most heat tolerant sorghum cultivars did not offer much resilience to warming temperatures, with a temperature threshold of 33°C, beyond which sorghum yields start to decline^8^. Under natural conditions, especially at the tops of canopies, direct sunlight can be very intense and thus oversaturate the photosynthetic mechanism in C_4_ plants. Sorghum leaves had reduced stomatal conductance and net CO_2_ assimilation rates after 4 h exposure to HL mimicking nature sunlight^9^. To improve C_4_ crop yields, it is crucial to holistically approach how C_4_ plants respond to HL or HT, two of the most influential environmental factors that can compromise C_4_ photosynthesis.

HL responses have been studied extensively in C_3_ plants^10–15^. To cope with reactive oxygen species (ROS) production and photooxidative stress resulting from HL, C_3_ plants have evolved many protective mechanisms which act on different timescales^10,14^. Non-photochemical quenching (NPQ), especially its predominant component, energy-dependent quenching (qE), acts within seconds to dissipate excess light energy as heat^10,16^. The formation of qE depends on the thylakoid lumen pH, the photosystem II (PSII) polypeptide PsbS, and the accumulation of the xanthophyll pigment zeaxanthin^17–19^. In C_3_ plants, under HL, violaxanthin is converted to the intermediate pigment antheraxanthin which is then converted to zeaxanthin by the enzyme violaxanthin de-epoxidase^20^. Accumulation of zeaxanthin is also necessary for induction of a slower-relaxing component of NPQ, zeaxanthin-dependent quenching (qZ)^21^. State transitions, which restructure the light harvesting complexes (LHCs) around PSII and PSI, occur on the order of minutes^10,16^. When photoprotective processes are insufficient, HL can result in photoinhibition (qI), which takes hours to recover^10^. Following HL exposure, expansion of the thylakoid lumen, swelling of the grana margin, and de-stacking of the thylakoid grana facilitate PSII repair by promoting accessibility and repair of PSII machinery^15,22–24^.

HL stress also results in dynamic transcriptional regulation of photosynthetic genes and induces the abscisic acid (ABA) pathway in Arabidopsis^11^.

HT is known to affect many cellular processes in C_3_ plants, including various aspects of photosynthesis^25–29^. C_3_ plants under HT have shown decreases in photosynthetic rates, inactivation of Rubisco, reduction of plastoquinone (PQ), and increase in cyclic electron flow (CEF) around photosystem I (PSI)^30^. Arabidopsis leaves treated with HT of 40°C has swollen M chloroplasts and increased plastoglobuli (PG) formation^31^. PG are thylakoid-associated plastid lipoprotein particles whose size, shape, and counts respond to abiotic stresses^32^. Additionally, HT induces the expression of heat shock transcription factors (HSFs), many of which have been implicated in transcriptional responses to numerous abiotic stresses, including HL and HT^33^. The induced HSFs increase the expression of heat shock proteins (HSPs), which are chaperone proteins involved in proper protein folding in response to HT and other abiotic stresses^34^.

Unlike C_3_ plants, studies on how C_4_ plants respond to HL or HT are largely limited, especially the underlying coordination between M and BS cells and the multi-level effects of HL and HT on photosynthesis, transcriptomes, and ultrastructure of C_4_ plants. A recent study examined the effects of HL stress in the C_4_ grass *Setaria viridis* over four days, with sampling points for photosynthetic parameters, sugar quantification, and transcriptome analyses every 24 hours^35^. They reported relatively minor transcriptional changes but a large accumulation of sugars without repression of photosynthesis in HL-treated samples^35^. These results suggest that prolonged HL-treated leaves undergo adaptive acclimation and transcriptional homeostasis in a few days. However, the short-term transcriptional responses of C_4_ plants to HL remain largely unknown. In sorghum leaves, HL induced the avoidance response in M chloroplasts and the swelling of BS chloroplasts (by cross section light microscope images), but the underlying mechanisms are unclear^9^. Research about how C_4_ photosynthesis responds to HT is mainly limited to biochemical and gas exchange analyses which suggest that HT results in Rubisco activation^36^, affects the activities of C_4_ carbon fixation enzymes^37^, decreases the BS conductance while increases CO_2_ leakiness^38,39^. Two transcriptome analyses in maize under HT have been reported^40,41^, but thorough analysis of C_4_ transcriptome with multi-level effects under HT is rare. Additionally, ultrastructural analysis in C_4_ plants under HL or HT can help us understand how HL or HT limits C_4_ photosynthesis and affects the coordination between M and BS cells, but currently such information is lacking.

To gain deeper insights into the molecular and physiological responses of C_4_ plants to HL or HT, we used the green foxtail *Setaria viridis* as a model. *S. viridis* is an excellent model to study C_4_ photosynthesis because of its expanding genetics and genomics toolkit, common growth condition, relatively quick generation time (8∼10 weeks, seed to seed), and similarity to important agronomic C_4_ crops, e.g. maize and sorghum^2,42,43^. We hypothesized that HL or HT affected C_4_ plants at different levels and linking multi-level changes could improve our understanding of HL or HT tolerance in C_4_ plants. We investigated the response of *S. viridis* to moderately HL or HT over a four-hour time-course at photosynthetic, ultrastructural, and transcriptomic levels (Fig. 1a). We monitored the dynamic changes of transcriptomes, pigments, and ABA levels over 4 h time points during the different treatments. We also measured photosynthetic parameters and ultrastructural changes after 4 h treatments, which revealed cumulative changes associated with the different treatments.

**Figure 1:**
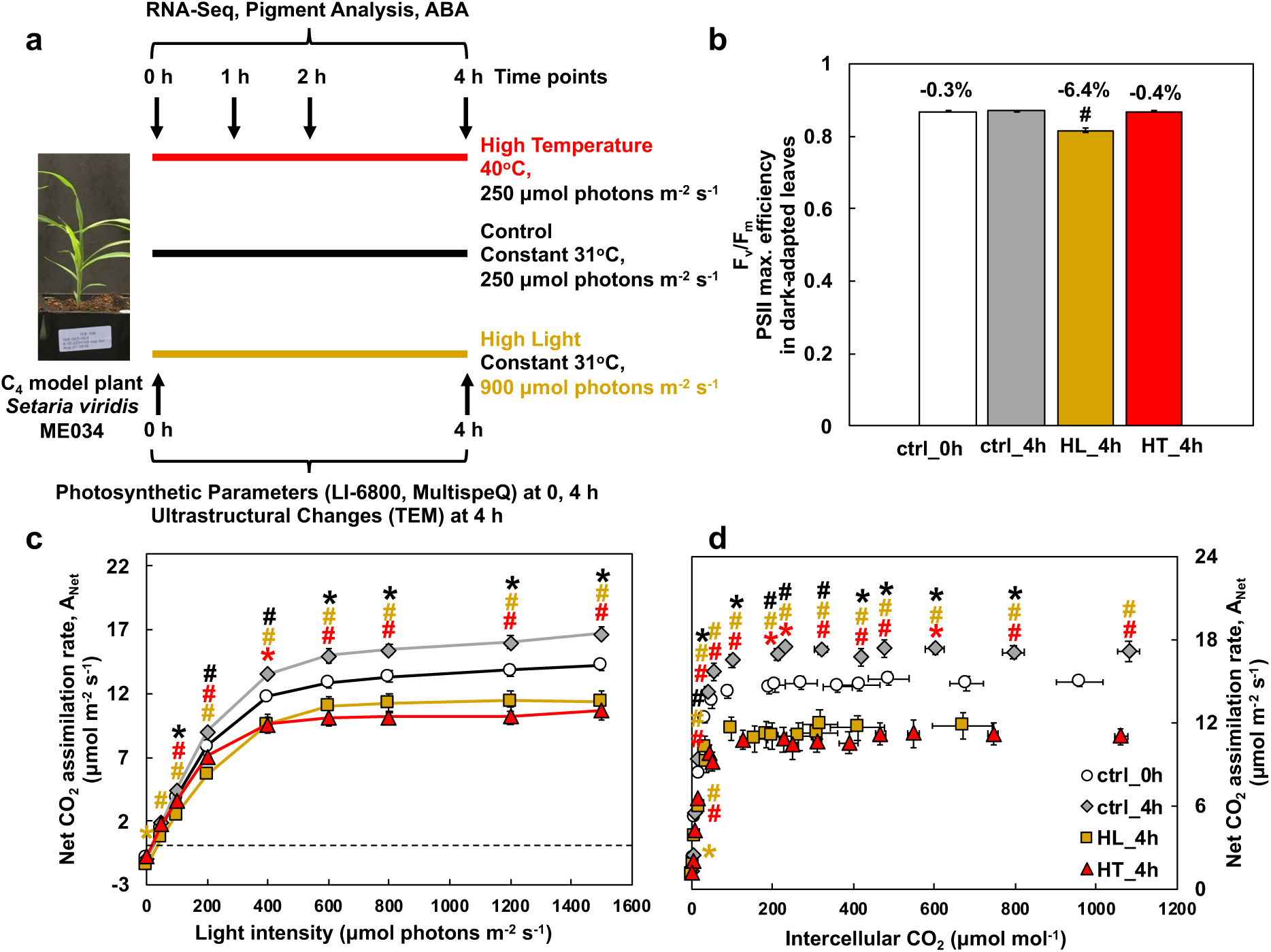
High light (HL) and high temperature (HT) resulted in comparable reduction of net CO_2_ assimilation rates and HL also caused significant photoinhibition in *S. viridis* leaves. **(a)** Experimental overview. We investigated how the C_4_ model plant *S. viridis* ME034 responded to HL or HT at different levels. Plants were grown under the control condition (31°C and 250 μmol m^−2^ s^−1^ light) for 13 days, then treated with control growth condition or HL (31°C, 900 μmol m^−2^ s^−1^) or HT (40°C, 250 μmol m^−2^ s^−1^ light) in different growth chambers for 4 h. The fourth fully expanded true leaves were utilized for all analyses. Leaf tissues from different treatments were harvested at 0, 1, 2, and 4 h time points for the analysis of RNA-seq, pigments, and leaf ABA levels. Photosynthetic parameters were measured using intact leaves at 0 and 4 h time points, including gas exchange and chlorophyll fluorescence using LI-6800 and spectroscopic measurements using MultispeQ. Transmission Electron Microscopy (TEM) analysis was performed to investigate chloroplast ultrastructure changes in leaves after 4 h treatments. **(b)** HL-treated leaves had reduced PSII maximum efficiency (F_v_/F_m_). F_v_/F_m_ was measured by chlorophyll fluorescence in LI-6800 with 20 min dark-adapted leaves. Pound symbols indicate statistically significant differences of ctrl_0h (at the start of treatments), HL_4h (after 4 h HL), and HT_4h (after 4 h HT) compared to ctrl_4h (after 4 h control treatment) using Student’s two-tailed t-test with unequal variance (# p<0.01). Percentages indicate reduction in F_v_/F_m_ compared to ctrl_4h. **(c, d)** Net CO_2_ assimilation rates during light response and CO_2_ response, respectively. Asterisk and pound symbols indicate statistically significant differences of ctrl_0h, HL_4h, and HT_4h compared to ctrl_4h using Student’s two-tailed t-test with unequal variance. P-values were corrected for multiple comparisons using FDR (*0.01<p<0.05, #p<0.01, the colors of * and # match the significance of the indicated conditions, black for ctrl_0h, yellow for HL_4h, red for HT_4h). Mean ± SE, *n* = 3-6 biological replicates.

Although we observed a comparable reduction of photosynthetic efficiency in HL- or HT-treated leaves, detailed analysis at multiple levels revealed different acclimation strategies to these two stresses in *S. viridis*. The transcriptional changes under HT were much less but more dynamic than under HL. The HL-treated leaves had over-accumulated starch in both M and BS chloroplasts, which may increase chloroplast crowdedness and inhibit PSII repair. While both HL and HT induced PG formation in chloroplasts, HT-treated leaves also had swollen chloroplasts and grana in M cells. Additionally, we observed different responses of M and BS cells under HT or HL and demonstrated the crosstalk between HL response and ABA signaling in C_4_ plants. Our research provides a systematic analysis of HL and HT responses in *S. viridis* and identifies potential targets to improve stress tolerance in C_4_ crops.

## Results

### HL or HT caused comparable reduction of photosynthesis and HL also resulted in photoinhibition

*S. viridis* leaves treated with 4 h HL (HL_4h) exhibited significantly reduced maximum efficiencies of PSII (F_v_/F_m_) as compared to that in 4 h control leaves (ctrl_4h) (Fig. 1b), suggesting HL-induced photoinhibition. Net CO_2_ assimilation rates (A_Net_) were significantly reduced in HL or HT-treated leaves in response to changes in light or CO_2_ (Fig. 1c,d). Pre-treatment control leaves (ctrl_0h) also had lower A_Net_ as compared to ctrl_4h leaves, suggesting circadian regulation of photosynthesis over the course of the day. The comparisons between different treatments at the 4 h time point should exclude the effects of circadian regulation. Stomatal conductance and transpiration rates in response to light were reduced in HL_4h leaves, especially at the beginning of the light response curve (Supplementary Fig. 3a,c). Stomatal conductance and transpiration rate in response to CO_2_ were lower in HL_4h or HT_4h leaves than ctrl_4h leaves (Supplementary Fig. 3b,d). PSII efficiency and electron transport rates in light-adapted leaves were reduced in HL_4h leaves as compared to ctrl_4h leaves in response to light (Supplementary Fig. 3e, g).

To estimate and model a variety of photosynthetic parameters, we assessed various aspects of leaf-level gas exchange measurements based on the light response curves and CO_2_ response curves (Supplementary Fig. 4). HL or HT compromised photosynthetic capacities and reduced several photosynthetic parameters in HL_4h and HT_4h leaves compared to ctrl_4h leaves, including gross maximum CO_2_ assimilation rates (*A_max_*), maximum carboxylation rates (*V_cmax_*), and quantum yields of CO_2_ assimilation (Supplementary Fig. 4a,b,c). HL_4h leaves had reduced stomatal conductance (*g_s_*) but increased light compensation point as compared to ctrl_4h leaves (Supplementary Fig. 4e, g). HT_4h leaves had reduced light saturation point as compared to ctrl_4h leaves (Supplementary Fig. 4h).

### Transcriptomics revealed important differences in the key pathways responding to HL or HT

To investigate the transcriptional patterns that may underlie the photosynthesis phenomena observed above, we performed RNA-seq analysis (Fig. 1a). Principal component analysis (PCA) of transcripts per million (TPM) (Supplementary Data 1) normalized read counts from ctrl, HL, and HT treatments showed the experimental conditions dominated the variance in the dataset (Fig. 2a).

**Figure 2:**
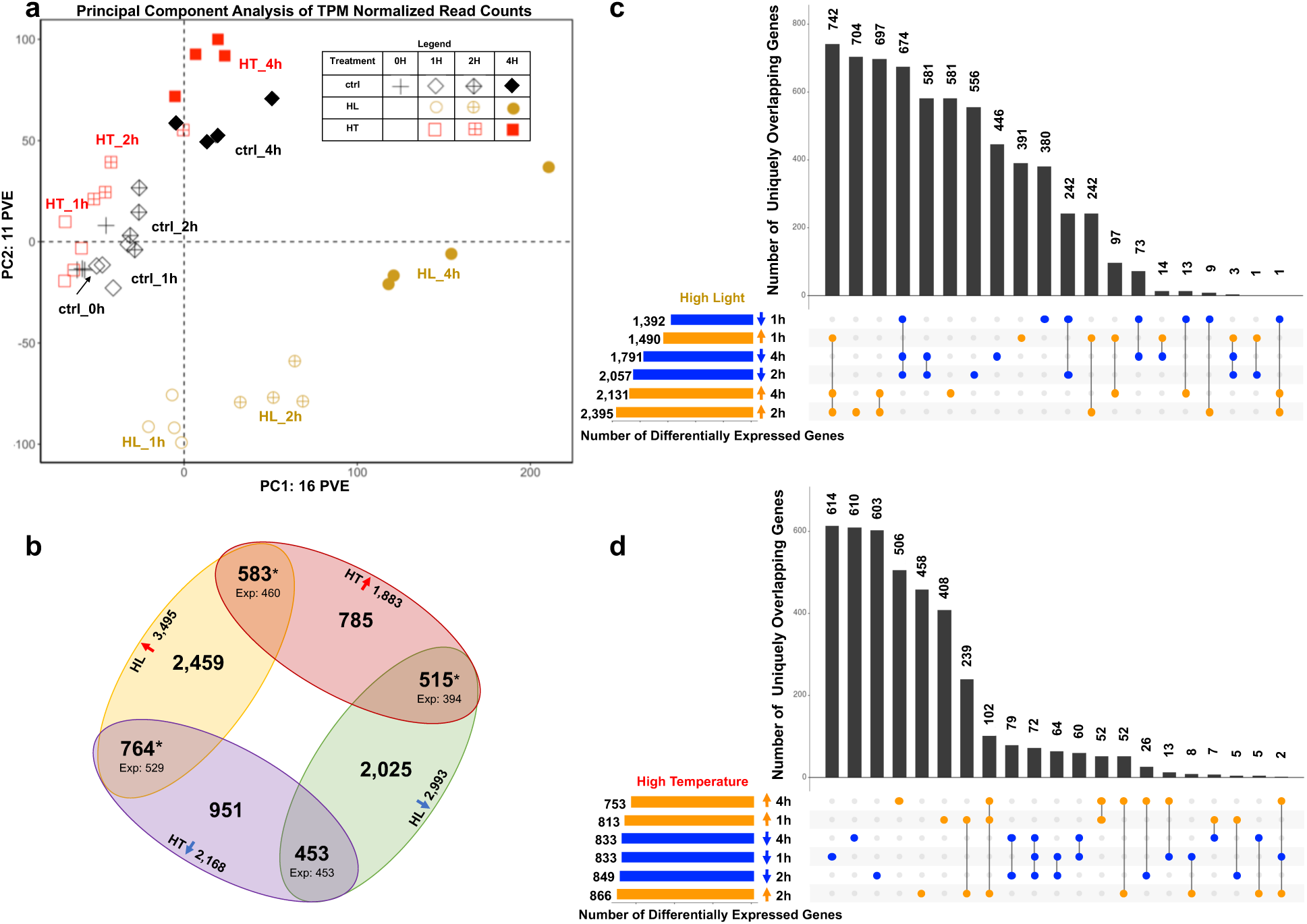
Time course transcriptome data reveal dynamic responses to high light or high temperature stresses in *S. viridis*. **(a)** Principal Component Analysis of TPM (transcripts per million) normalized read counts in control (ctrl), high light (HL), and high temperature (HT) treated samples. The first two principal components representing the highest percent variance explained are displayed. PC1 explains 16% of the variance in the dataset and mainly separates the samples based on time. PC2 explains 11% of the variance in the dataset and mainly separates the HL samples from the ctrl and HT samples. Black diamonds indicate ctrl samples, yellow circles indicate HL samples, and red squares indicate HT samples. Different fillings for these symbols indicate different time points of each treatment. Each treatment and time point have four biological replicates, represented by symbols with the same shape and color. **(b)** HL and HT treatments had more overlapping differentially expressed genes than expected by random chance. Gene sets represent the number of genes differentially regulated in at least one time point in the given condition. Red upward arrows denote up-regulation and blue downward arrows denote down-regulation. Yellow oval denotes HL up-regulated genes, green oval denotes HL down-regulated genes, red oval denotes HT up-regulated genes, purple oval denotes HT down-regulated genes. Expected values (Exp) are the number of the overlapping genes expected by random chance based on size of the gene lists and background of all genes tested via DeSeq2 (14,302). Numbers above expected values are the actual number of overlapped genes between two conditions. *p<0.0001, Fisher’s Exact Test. **(c, d)** HT transcriptional responses are more transient than HL. UpSetR plots show number of uniquely overlapping genes between up and down regulated gene sets at each time point in HL and HT, respectively. Horizontal bars indicate the number of genes up or down regulated at each time point. Filled circles indicate the gene sets included in the overlap shown. Vertical bars indicate the number of genes represented in the overlap shown. Overlapping gene sets are arranged in descending order by number of genes. Genes may only belong to a single overlapping gene set and are sorted into the overlapping set with the highest number of interactions.

Next, we compared differentially expressed genes (DEGs) between HL and HT treatments. Genes that were either up- or down-regulated in at least one time point were included in the lists of DEGs for each condition. Utilizing this method, we were able to broadly compare the trends between the HL and HT transcriptomes. There were more DEGs identified in the HL dataset than the HT dataset (Fig. 2b, Supplementary Fig. 5 and Data 2). Significantly more genes were up-regulated in both HL and HT-treated transcriptomes than would be expected by random chance (Fig. 2b, Supplementary Data 4). Additionally, significantly more genes were regulated in opposite directions between HL and HT transcriptomes than would be expected by random chance. To visualize how DEGs were conserved between time points within treatments, we plotted the overlaps between up- and down-regulated genes at each time point. In HL-treated samples, 742 genes were up-regulated at 1, 2, 4 h time points, representing the largest subset of uniquely overlapping genes and the core HL-induced genes (Fig. 2c, Supplementary Data 3). Similarly, 674 genes were down-regulated at all three time points of HL treatment, representing the core HL-reduced genes. Conversely, in the HT-treated samples, the expression pattern was dominated by genes differentially expressed at a single time point (Fig. 2d), indicating the transcriptional response to HT was more transient and dynamic than that to HL. In HT-treated samples, 102 and 72 genes were up- and down-regulated at all three time points, representing the core HT induced and reduced genes, respectively.

To reveal transcriptional changes that may explain the reduced photosynthesis under HL or HT, we grouped DEGs into several key pathways. Investigation of genes related to the light reaction of photosynthesis revealed that many genes involved in PSII assembly/repair and photoprotection (e.g., *PsbS*), were up-regulated in HL, while many genes relating to LHCII and the core complexes of PSII/PSI were down regulated in HL (Fig. 3a,b). Although HT treatment did not result in the same extent of differential regulation of light reaction related genes as HL, STN7, a kinase involved in state 1 to state 2 transitions^44^ was induced, while TAP38, a phosphatase involved in state 2 to state 1 transitions^45^, was repressed in HT-treated leaves (Supplementary Fig. 6a). This suggests a possible heat induced state transition to move the mobile LHCII from PSII (state 1) to PSI (state 2). Additionally, several genes related to the chloroplast NDH (NADPH dehydrogenase) complex were up-regulated in the HT treatment (Fig. 3b). Furthermore, when investigating genes involved in cyclic electron flow (CEF) (Supplementary Fig. 6), we found that key components of CEF, *PGR5 (proton gradient regulation 5)*^46^ and two copies of *PGRL1 (PGR5-like photosynthetic phenotype 1)*^47^, were induced under HT, suggesting heat-induced CEF around PSI.

**Figure 3:**
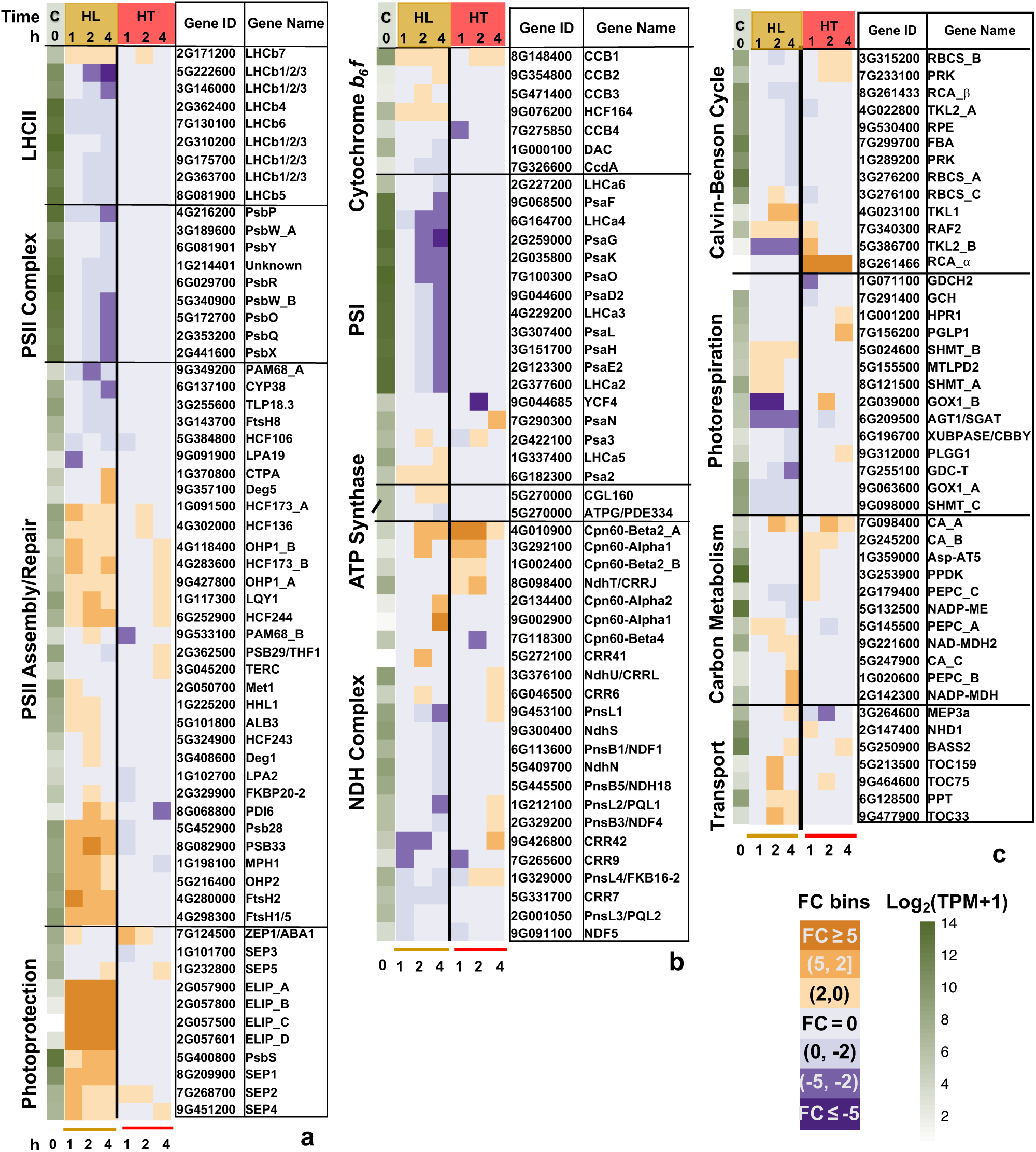
High light (HL) differentially regulated genes involved in photosynthesis more than high temperature (HT). **(a, b)** Genes related to light reaction of photosynthesis and photoprotection. **(c)** Genes related to carbon metabolism and chloroplast transport. The first green column displays log_2_(mean TPM + 1) at ctrl_0h (at the start of treatments, C). TPM, transcripts per million, normalized read counts. Heatmap displays the fold change (FC) bin of DeSeq2 model output values at 1, 2, 4 h of HL or HT versus control at the same timepoint (q < 0.05). FC bins: highly induced: FC ≥ 5; moderately induced: 5 > FC ≥ 2; slightly induced: 2 > FC > 0; not differentially expressed: FC = 0; slightly repressed: 0 > FC > −2; moderately repressed: −2 ≥ FC > −5; highly repressed: FC ≤ −5. Gene ID: *S. viridis* v2.1 gene ID, excluding “Sevir.”. All genes presented in the heatmaps were significantly differentially regulated in at least one time point.

Under HL treatment, the transcriptional changes of genes involved in the Calvin-Benson cycle were less than those involved in the light reactions of photosynthesis (Fig. 3c). Rubisco activase (RCA) is essential for CO_2_ fixation by maintaining the active status of Rubisco^48,49^. The *S. viridis* genome has two adjacent genes encoding RCAs (Supplementary Fig. 7). Protein sequence alignment of the two *S. viridis* RCAs with Arabidopsis RCAs revealed one SvRCA-α which retains the two conserved redox-sensitive cysteine residues as in AtRCA*_*α, and one *SvRCA_*β which has higher basal expression (approximately 700-fold higher) than Sv*RCA_*α and possibly the major RCA in *S. viridis*. *SvRCA_*α was highly induced during the entire 4 h HT (Fig. 3c).

Key genes involved in photorespiration, e.g. *GOX1* (glycolate oxidase)^50,51^ and *AGT1* (Serine:glyoxylate aminotransferase)^52^ were down-regulated under HL (Fig. 3c). *GOX1* and several other genes involved in photorespiration, *PGLP1* (2-phosphoglycolate phosphatase)^53^, *HPR1* (hydroxypyruvate reductase)^54^, and *PLGG1* (plastidic glycolate/glycerate transporter)^55^ were induced under HT, suggesting heat-induced photorespiration.

Some genes important for C_4_ carbon metabolism were up-regulated under HL (Fig. 3c), e.g. *PEPC_B* (phosphoeynylpyruvate carboxylase) and *NADP-MDH* (NAD-dependent malate dehydrogenase)^1^. Carbonic anhydrase^56^ (CA_A) was induced under both HL and HT.

By investigating pathways associated with photosynthesis, we found HL increased the expression of starch biosynthesis/degradation genes and genes encoding PG-localized proteins (Fig. 4a), but down-regulated several genes in the sugar-sensing pathway (Fig. 4b), and differentially regulated several sugar transporter genes (Supplementary Fig. 6b). These transcriptional changes were much less pronounced under HT.

**Figure 4:**
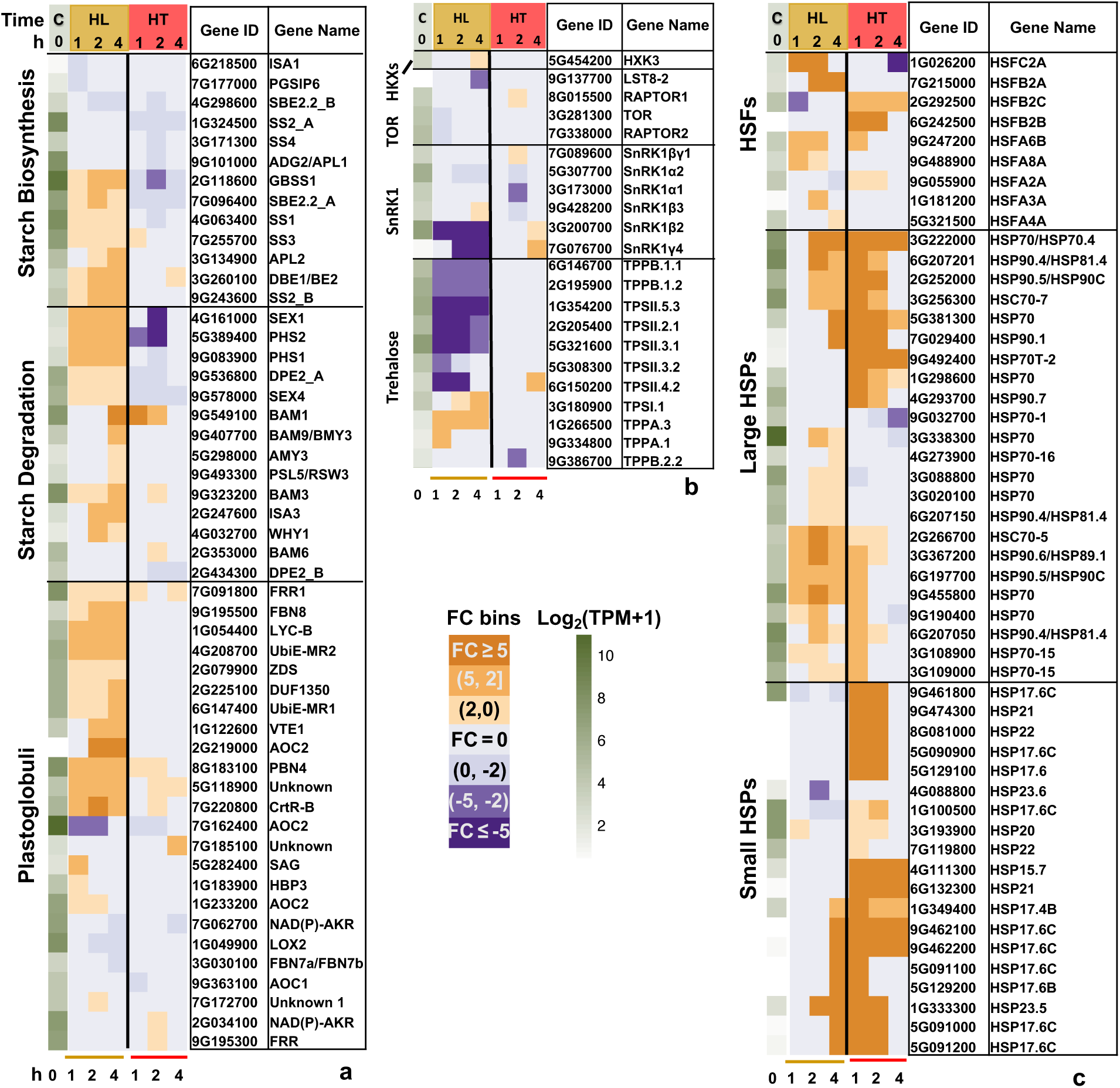
High light (HL) and high temperature (HT) differentially regulated genes involved in several key pathways. **(a, b)** HL induced genes involved in starch biosynthesis/degradation and genes encoding plastoglobuli-localized proteins; **(b)** HL down-regulated many genes of the sugar-sensing pathways; **(c)** Both HL and HT induced genes encoding shock transcription factors (HSFs) and heat shock proteins (HSPs) but the induction was much quicker under HT than HL. The first green column displays log_2_(mean TPM + 1) at ctrl_0h (at the start of treatments, C). TPM, transcripts per million, normalized read counts. Heatmap displays the fold change (FC) bin of DeSeq2 model output values at 1, 2, 4 h of HL or HT versus control at the same timepoint (q < 0.05). FC bins: highly induced: FC ≥ 5; moderately induced: 5 > FC ≥ 2; slightly induced: 2 > FC > 0; not differentially expressed: FC = 0; slightly repressed: 0 > FC > −2; moderately repressed: −2 ≥ FC > −5; highly repressed: FC ≤ −5. Gene ID: *S. viridis* v2.1 gene ID, excluding “Sevir.”. All genes presented in the heatmaps were significantly differentially regulated in at least one time point.

Several HSFs had highly induced expression under either HL or HT, but interestingly, different HSFs were up-regulated in HL vs HT (Fig. 4c). *HSFA6B* was a notable exception, which was induced in both HL and HT. A set of shared HSPs were induced under both HT and HL, but the induction was quicker and stronger under HT than HL, especially the small HSPs, suggesting shared but also temporally distinct transcriptional responses of HSPs under HL and HT.

We also investigated genes associated with ROS pathways. Specialized ROS scavenging pathways have evolved in plants^57^. We identified genes encoding antioxidant enzymes in *S. viridis* and investigated their expression patterns under HL or HT (Supplementary Fig. 6c). Three gene families of antioxidant enzymes have many members with strong differential expression in HL-treated leaves: TRX (thioredoxin), POX (peroxidases), and GST (glutathione S-transferase). Interestingly, within each of the three antioxidant pathways, some genes were up-regulated while others were down-regulated in HL-treated leaves. A similar pattern was shown in HT-treated leaves, although with fewer differentially regulated genes.

The reduced stomatal conductance in HL_4h leaves (Supplementary Fig. 3a) suggested there may be changes in ABA pathways and leaf ABA levels. Our RNA-seq analysis showed that several genes in the ABA pathways were up-regulated in response to HL (Fig. 5a). Additional, ABA levels were increased 3-fold in HL_1h leaves followed by a return to baseline by HL_4h (Fig. 5b).

**Figure 5:**
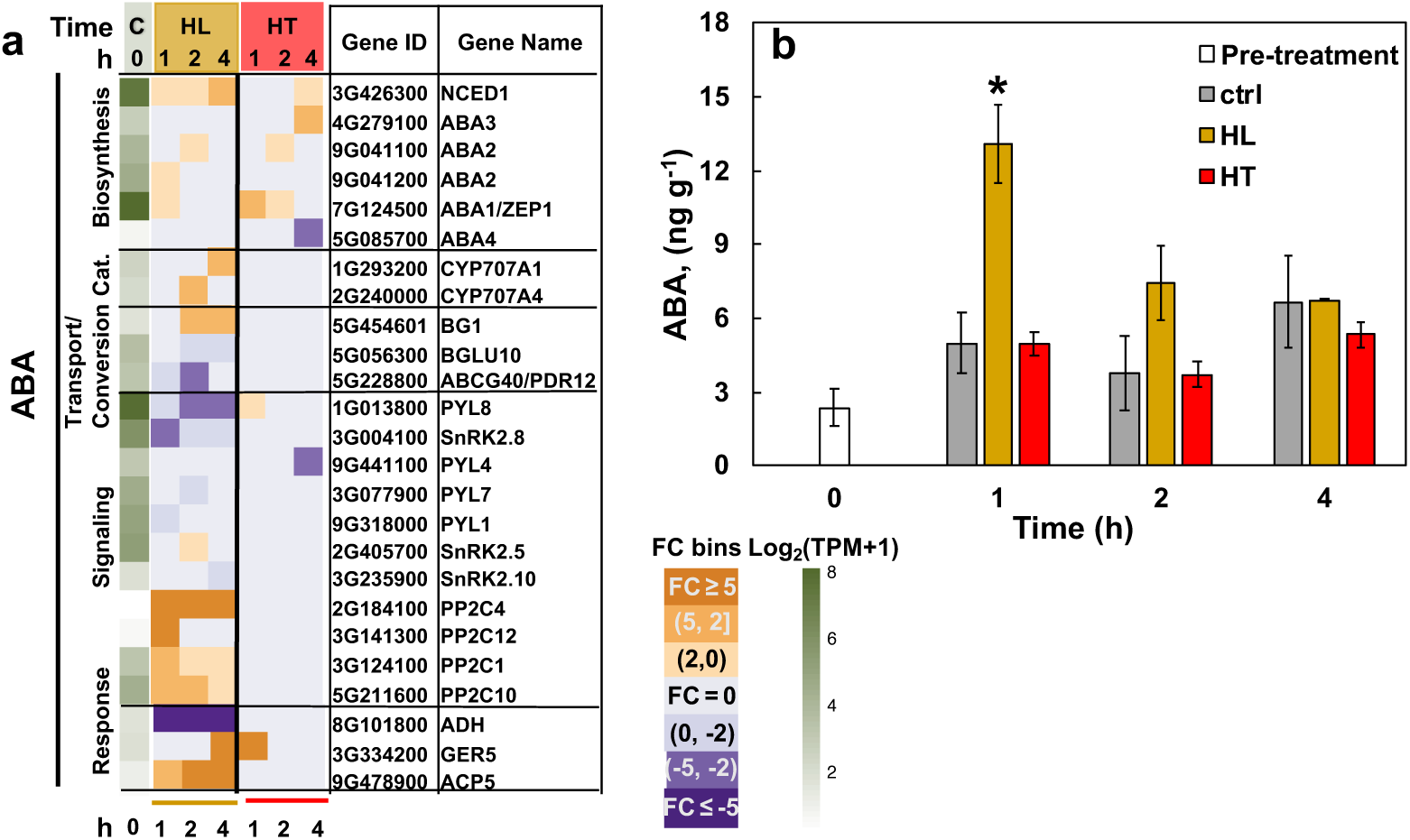
High light (HL) up-regulated genes involved in the abscisic acid (ABA) pathway and transiently increased leaf ABA levels. **(a)** Heatmap of differentially regulated genes involved in the ABA pathway. Cat: catabolism. The first green column displays log_2_(mean TPM + 1) at ctrl_0h (at the start of treatments, C). TPM, transcripts per million, normalized read counts. Heatmap displays the fold change (FC) bin of DeSeq2 model output values at 1, 2, 4 h of HL or HT versus control at the same timepoint (q < 0.05). FC bins: highly induced: FC ≥ 5; moderately induced: 5 > FC ≥ 2; slightly induced: 2 > FC > 0; not differentially expressed: FC = 0; slightly repressed: 0 > FC > −2; moderately repressed: −2 ≥ FC > −5; highly repressed: FC ≤ −5. Gene ID: *S. viridis* v2.1 gene ID, excluding “Sevir.”. All genes presented in the heatmaps were significantly differentially regulated in at least one time point. **(b)** Concentrations of leaf ABA. Mean ± SE, *n* = 3 biological replicates. Asterisk symbol indicates statistically significant differences as compared to the control condition at the same time point. (Student’s two-tailed t-test with unequal variance, *0.01<p<0.05).

To distinguish M and BS specific transcriptomic responses and gain more information about how these two specialized cell types function together in HL or HT responses, we investigated the cell type specificity of our pathways of interest (Supplementary Fig. 8, Supplementary data 6) by using previously published M and BS specific transcriptomes under control conditions^58^. We observed several cell-type specific transcriptional responses to HL or HT, e.g. pathways related to ROS-scavenging, sugar transport, and HSPs.

### HL treatment induced NPQ in *S. viridis*

The increased photoinhibition in HL_4h leaves and the increased *PsbS* transcription under HL prompted us to quantify NPQ and xanthophyll pigments. NPQ was significantly higher in HL_4h leaves than ctrl_4h leaves in response to light and CO_2_ (Fig. 6a,b). The HL-induced NPQ measured by LI_6800 was confirmed using MultispeQ with the estimated NPQ, NPQ_(T),_ based on a method that estimates NPQ in light-adapted leaves^59^ (Supplementary Fig. 9c). The increased NPQ was also supported by the observed 4-fold increase of zeaxanthin (Fig 6c) during HL. Additionally, HL treatment doubled the intermediate antheraxanthin level (Fig. 6d) and tripled the overall de-epoxidation state of the xanthophyll cycle (Fig. 6e). In Arabidopsis, lutein also has a role in NPQ or qE and can substitute for zeaxanthin in qE formation^60^. Lutein as well as total carotenoids were significantly induced in HL_4h leaves (Supplementary Fig. 10c,d). These results indicate the occurrence of photoprotection in HL-treated leaves. Ctrl and HT treatments had little effect on leaf pigments.

**Figure 6:**
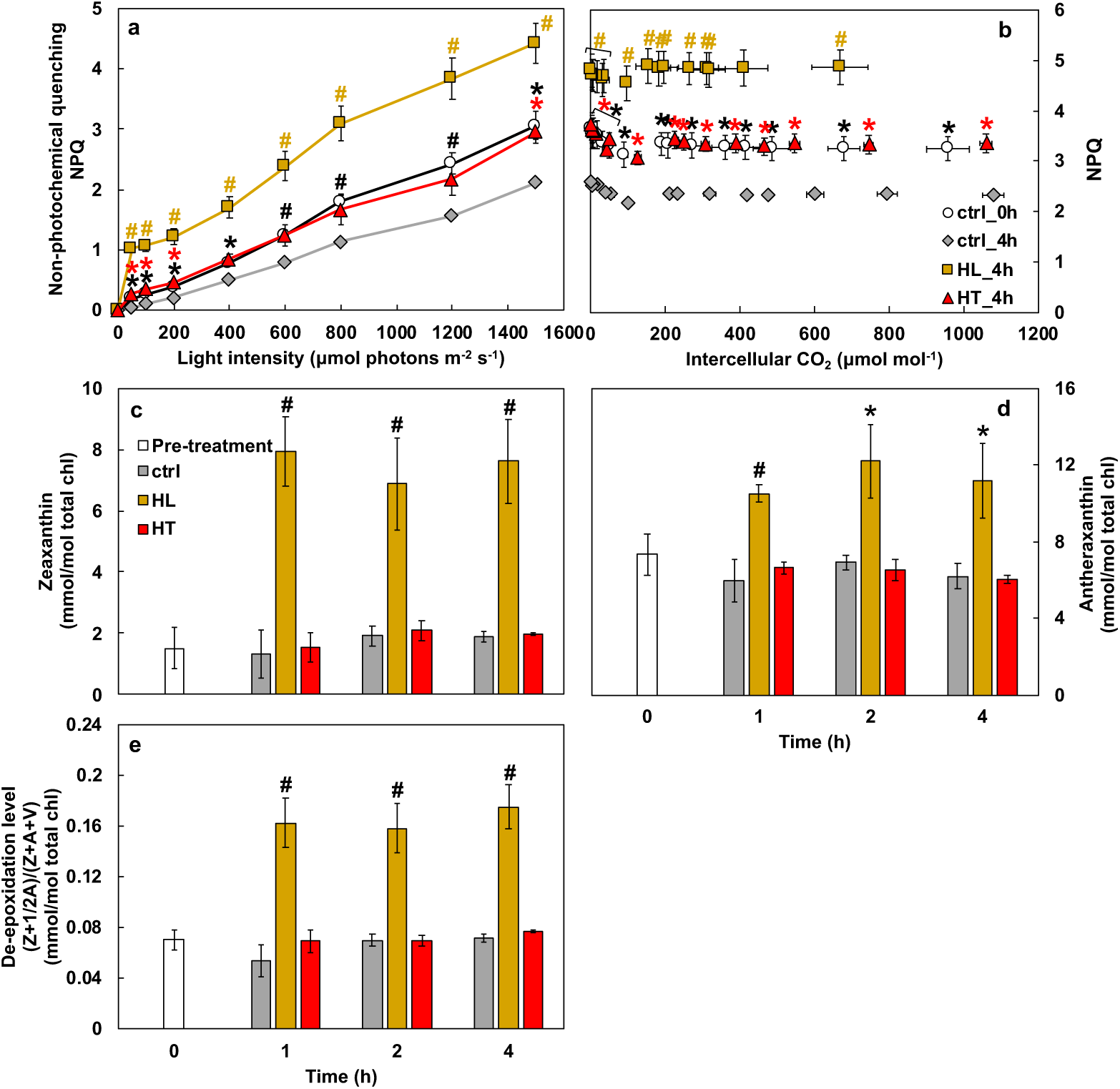
High light (HL) induced non-photochemical quenching (NPQ) and increased zeaxanthin as well as de-epoxidation levels. **(a)** Light and **(b)** CO_2_ response of NPQ. Mean ± SE, *n* = 3-6 biological replicates. Asterisk and pound symbols indicate statistically significant differences of ctrl_0h (at the start of treatments), HL_4h (after 4 h high light), and HT_4h (after 4 h high temperature) compared to ctrl_4h (after 4 h control treatment) using Student’s two-tailed t-test with unequal variance. P-values were corrected for multiple comparisons using FDR (*0.01<p<0.05, #p< 0.01, the colors of * and # match the significance of the indicated conditions, black for ctrl_0h, yellow for HL_4h, red for HT_4h). **(c, d, e)** Concentrations of zeaxanthin, antheraxanthin, and xanthophyll cycle de-epoxidation. Mean ± SE, *n* = 3 biological replicates. Asterisk and pound symbols indicate statistically significant differences of high light (HL) or high temperature (HT) treatments compared to the control (ctrl) condition at the same time points using Student’s two-tailed t-test with unequal variance (*0.01<p<0.05, #p<0.01).

### HL or HT altered chloroplast ultra-structures

The reduced photosynthesis (Fig. 1c,d) in HL_4h and HT_4h leaves, and the HL-induced photoinhibition (Fig. 1b) and transcripts related to the starch as well as PG pathways (Fig. 4a) led us to investigate the ultrastructural changes of the M and BS chloroplasts in ctrl_4h, HL_4h, and HT_4h leaves by using transmission electron microscopy (TEM).

TEM images showed HL_4h leaves had increased relative starch volume fraction and chloroplast area in both M and BS chloroplasts, but decreased relative volume fractions of stroma plus stroma lamellae (unstacked thylakoid membranes) in M chloroplasts as compared to ctrl_4h leaves (Fig. 7, Supplementary Fig. 15), suggesting increased starch accumulation and chloroplast crowdedness under HL. Starch quantification using biochemical assays confirmed 3x starch levels in HL_4h leaves as compared to ctrl_4h leaves (Fig. 7m). In HT_4h leaves, M chloroplasts had reduced relative starch volume fraction but increased chloroplast area (Fig. 7, Supplementary Fig. 15). HT did not affect the relative volume of stroma or stroma lamellae in either M or BS chloroplasts.

**Figure 7:**
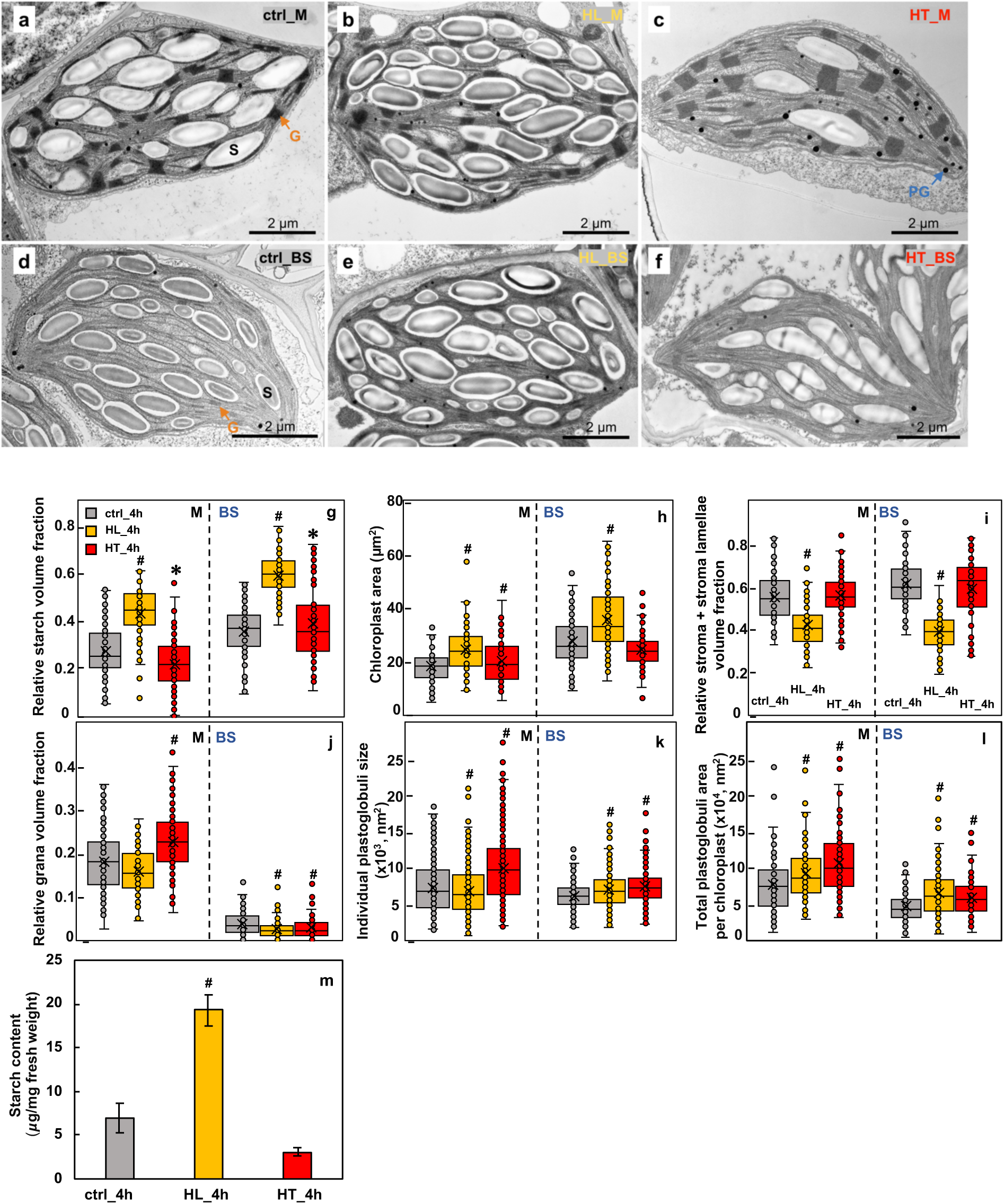
High light (HL) increased starch accumulation and both HL and high temperature (HT) treatments induced chloroplast plastoglobuli formation in *S. viridis* leaves. **(a-f)** Representative transmission electron microscopy (TEM) images of mesophyll (M) and bundle sheath (BS) chloroplasts in leaves of *S. viridis* after 4 h treatments of control (ctrl_4h, 31°C, 250 μmol m^−2^ s^−1^ light) or high light (HL_4h, 31°C, 900 μmol m^−2^ s^−1^) or high temperature (HT_4h, 40°C, 250 μmol m^−2^ s^−1^ light). TEM images of M **(a, b, c)** and BS **(d,e,f)** chloroplasts. S labels the starch granule; G labels grana, the orange arrows indicate grana in M and BS chloroplasts; PG labels plastoglobuli. **(g, i, j)** Relative volume fraction of indicated parameters were quantified using Stereo Analyzer with Kolmogorov–Smirnov test for statistical analysis compared to the same cell type of the control condition. **(h, k, l)** area and size of indicated parameters were quantified using ImageJ with two-tailed t-test with unequal variance compared to the same cell type of the control condition. Each treatment had three biological replicates, total 90-120 images per treatment. *0.05<p<0.01; #p<0.01. (**m**) Starch quantification using starch assay kits. HL_4h leaves accumulated 4x starch as compared to ctrl_4h leaves. Values are mean ± SE, *n* = 3 biological replicates. Pound symbols indicate statistically significant differences as compared to ctrl_4h using Student’s two-tailed t-test with unequal variance (#p< 0.01).

Like in other C_4_ plants, grana in *S. viridis* are dominantly present in the M chloroplasts. BS chloroplasts also have some grana, which are absent from the central area but present in the peripheral region (Fig. 7d-f). HL reduced grana width in M chloroplasts and the relative volume, height, and area of grana in BS chloroplasts as compared to the ctrl condition (Fig. 7j, Supplementary Fig. 14,15). The HT effects on grana structure were quite different from HL. M chloroplasts at HT had increased relative volume, height, area, and mean layer thickness of grana, indicating heat-induced grana swelling. However, in BS chloroplasts, HT decreased the relative volume, width, and area of grana, suggesting that HT affected grana structure differently in M and BS chloroplasts.

HL increased PG count and the total PG area per chloroplast, while it decreased the mean individual PG size in M chloroplasts, indicating smaller but more numerous PGs in M chloroplasts (Fig. 7k, l, Supplementary Fig. 13). Furthermore, HL increased individual PG size and total PG area per chloroplast in BS chloroplasts (Supplementary Fig. 13). HT increased individual PG size and total PG area, suggesting heat-induced PG formation in both M and BS chloroplasts.

### HL- and HT-treated Leaves had reduced photosynthetic capacity

The over-accumulated starch in HL_4h leaves (Fig. 7) and the increased leaf ABA levels (Fig. 5) led us to investigate photosynthesis under the simulated stress conditions and immediately after different treatments without dark-adaptation in the LI_6800 leaf chamber (Fig. 8). Under the same temperature and light intensity in the LI-6800 leaf chamber, most photosynthetic parameters with or without dark-adaptation were similar (groups 1 vs. 2) (Fig. 8). Under the simulated treatment condition in the LI-6800 leaf chamber (group 3), HL_4h leaves had higher net CO_2_ assimilation rates (A_Net_) and stomatal conductance under 600 µmol photons m^−2^ s^−1^ light than ctrl_4h leaves under 200 µmol photons m^−2^ s^−1^ light, but both parameters in HL_4h leaves were lower than ctrl_4h leaves under the same light intensity (group 3, 4) (Fig. 8a). This suggests that HL_4h leaves had reduced capacities for A_Net_ and stomatal conductance as compared ctrl_4h leaves under the same condition. Under the simulated treatment condition (group 3), HL_4h leaves under 600 µmol photons m^−2^ s^−1^ light had reduced PSII operating efficiency (Fig. 8c), increased electron transport rates (Fig. 8d), and increased NPQ (Fig. 8f) as compared to the ctrl_4h leaves under 200 µmol photons m^−2^ s^−1^ light, consistent with light induced electron transport and NPQ.

**Figure 8:**
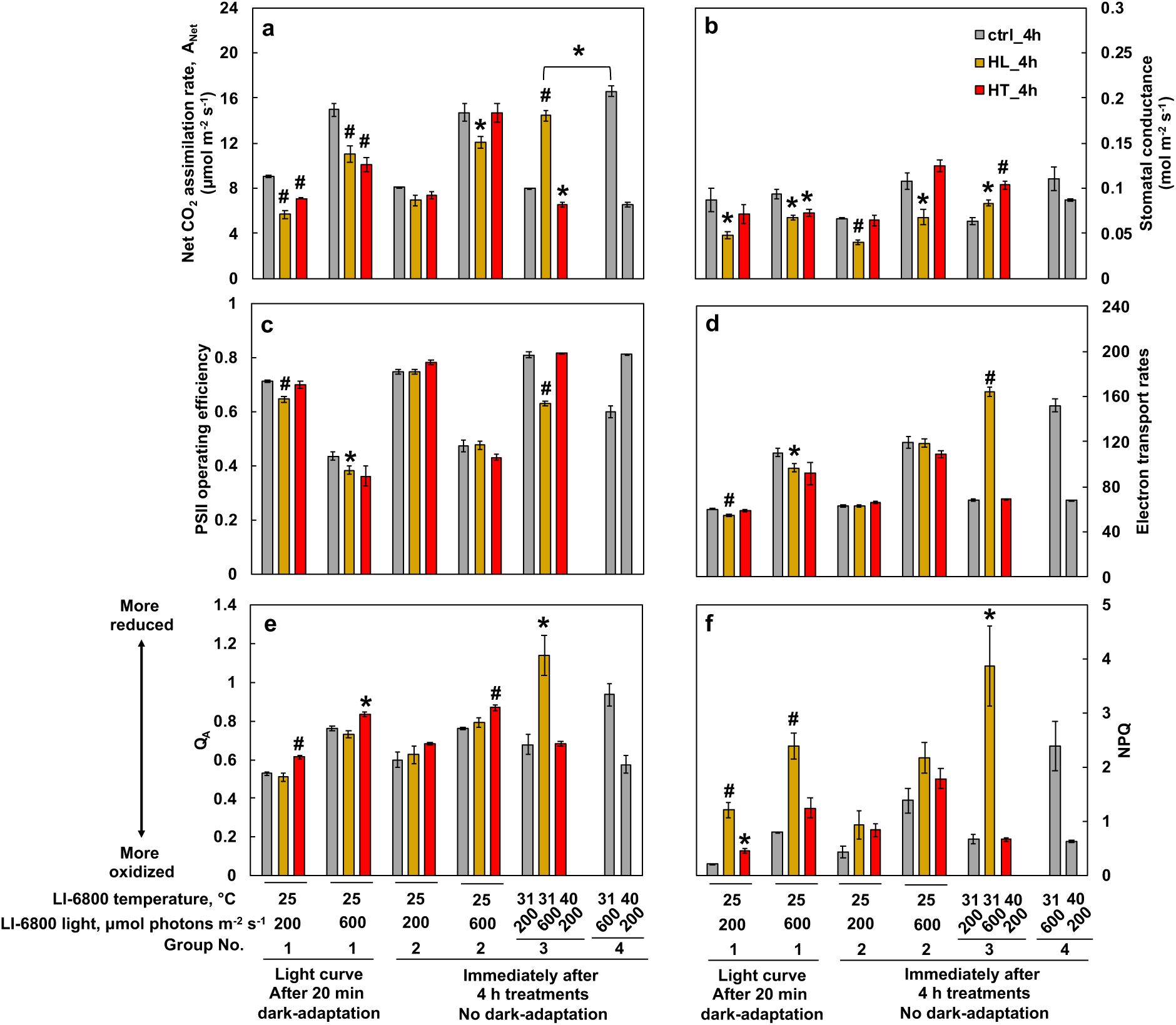
High light or high temperature treated leaves had lower photosynthetic capacities than leaves treated with the control condition. *S. viridis* plants were treated with 4 h of control growth condition (ctrl_4h, 31°C, 250 μmol m^−2^ s^−1^ light) or high light (HL_4h, 31°C, 900 μmol m^−2^ s^−1^) or high temperature (HT_4h, 40°C, 250 μmol m^−2^ s^−1^ light) in different growth chambers. After the treatments, an intact fourth fully expanded true leaf from each treated plant was clamped in the LI-6800 leaf chamber to measure various photosynthetic parameters. Group 1 are select data from the light response curves after 20 min dark-adaptation in the LI-6800 leaf chamber with indicated light and temperature. Groups 2,3,4 were measured immediately after 4 h of ctrl, HL, HT treatments without dark-adaptation and under the indicated temperature and light condition in the LI-6800 leaf chamber. Individual plants were used for each replicate. **(a)** Net CO_2_ assimilation rates. **(b)** Stomatal conductance. **(c)** PSII operating efficiency. **(d)** Electron transport rate. (**e**) Plastoquinone redox status (Q_A_). (**f**) NPQ, Non-photochemical quenching. Asterisk and pound symbols indicate statistically significant differences of HL_4h and HT_4h leaves compared to ctrl_4h leaves in the same group or under the same LI-6800 leaf chamber condition using Student’s two-tailed t-test with unequal variance (*0.01<p<0.05, #p<0.01).

Without dark-adaptation, HT_4h leaves had similar A_Net_ as ctrl_4h leaves (Fig. 8a, group 2). This may be due to the transient recovery of photosynthesis after switching the HT_4h leaves from 40°C in the growth chamber to 25°C in the LI_6800 leaf chamber for measurements. Under the same light intensity, HT_4h leaves had significantly lower A_Net_ (Fig. 8a) and more reduced plastoquinone (Fig. 8e) than ctrl_4h leaves. Under the simulated treatment condition in LI-6800 leaf chamber (group 3), HT_4h leaves had increased stomatal conductance (Fig. 8b) but reduced A_Net_ as compared to ctrl_4h leaves (Fig. 8a), consistent with transpiration cooling of leaf temperature (Supplementary Fig. 1) and reduced photosynthetic capacity in HT-treated leaves (Supplementary Fig. 4a-c).

### The activity of ATP synthase was inhibited in HL-treated leaves

Based on the HL-induced starch accumulation, we hypothesized that starch may inhibit photosynthesis through feedback regulation. We measured electrochromic shift (ECS) and chlorophyll fluorescence using MultispeQ^61^ to evaluate proton fluxes and the transthylakoid proton motive force (*pmf*) *in vivo*^62–64^. Different treatments did not significantly change *pmf* (Fig. 9a). HL_4h leaves had significantly reduced proton conductivity and lower proton flux rates as compared to ctrl_4h leaves (Fig. 9b,c), indicating reduced ATP synthase activity in HL-treated leaves. The MultispeQ NPQ_(T)_ data showed that the HL-induced NPQ was more sensitive to *pmf* than ctrl_4h leaves, with higher NPQ produced at a given level of proton motive force in HL_4h leaves than ctrl_4h leaves (Fig. 9d).

**Figure 9:**
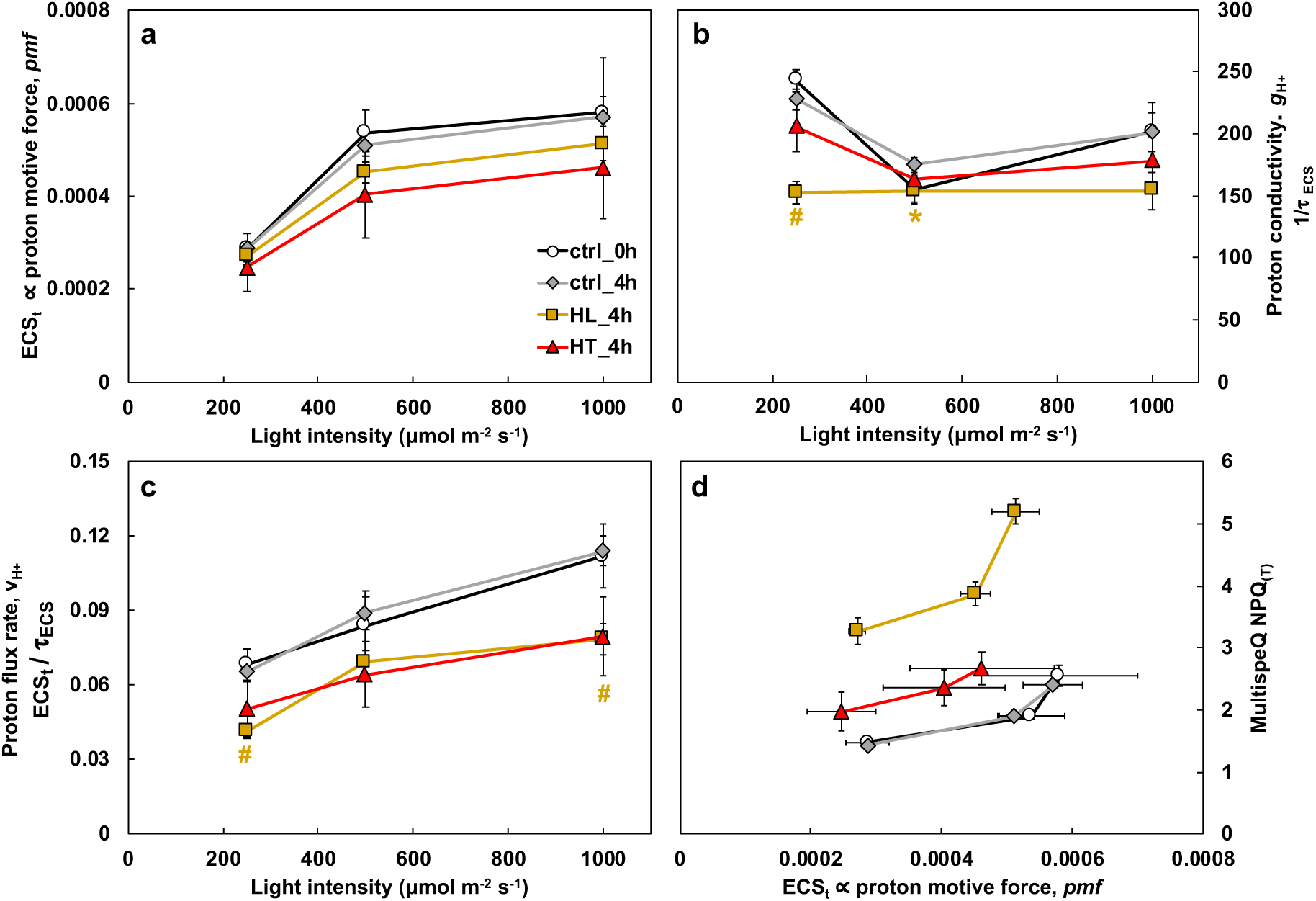
High light treatment inhibited ATP synthase activity. *S. viridis* plants were treated with 4 h of control growth condition (ctrl_4h, 31°C, 250 μmol m^−2^ s^−1^ light) or high light (HL_4h, 31°C, 900 μmol m^−2^ s^−1^) or high temperature (HT_4h, 40°C, 250 μmol m^−2^ s^−1^ light) in different growth chambers. After the treatments, photosynthetic parameters in treated leaves were monitored using the MultispeQ instrument. **(a)** ECS_t_, measured by electrochromic shift (ECS), representing the transthylakoid proton motive force, *pmf*. **(b)** Proton conductivity (ɡ_H_^+^ =1/_/_τ_ECS_), proton permeability of the thylakoid membrane and largely dependent on the activity of ATP synthase, inversely proportional to the decay time constant of light-dark transition induced ECS signal (τ_ECS_). **(c)** Proton flux rates, v_H+_, calculated by ECS_t /_τ_ECS_, the initial decay rate of the ECS signal during the light-dark transition and proportional to proton efflux through ATP synthase to make ATP. **(d)** Non-photochemical quenching (NPQ) measured by MultispeQ. Mean ± SE, *n* = 3 biological replicates. Asterisk and pound symbols indicate statistically significant differences of ctrl_0h, HL_4h, and HT_4h compared to ctrl_4h using Student’s two-tailed t-test with unequal variance. (*0.01<p<0.05, #p< 0.01, the colors of * and # match the significance of the indicated conditions, yellow for HL_4h).

## Discussion

We investigated how the C_4_ model plant *S. viridis* responds to HL or HT stresses at multiple levels by employing diverse approaches (Fig. 1a). Our data provide a thorough analysis of HL and HT responses in *S. viridis* at photosynthetic, transcriptomic, and ultrastructural levels and reveal limitations of photosynthesis under HL or HT. The HL (900 µmol photons m^−2^ s^−1^) and HT (40°C) treatments we chose were both moderate stresses within the physiological range for *S. viridis*. Although the impact of moderate stresses can be difficult to analyze due to mild phenotypes, moderate stresses are highly relevant and occur frequently in the field^65^. Understanding the impacts of moderate stresses on C_4_ plants is imperative for agricultural research. The moderately HL and HT we used reduced net CO_2_ assimilation rates at comparable levels in *S. viridis* leaves (Fig. 1c), but via different mechanisms (Fig. 10).

**Figure 10:**
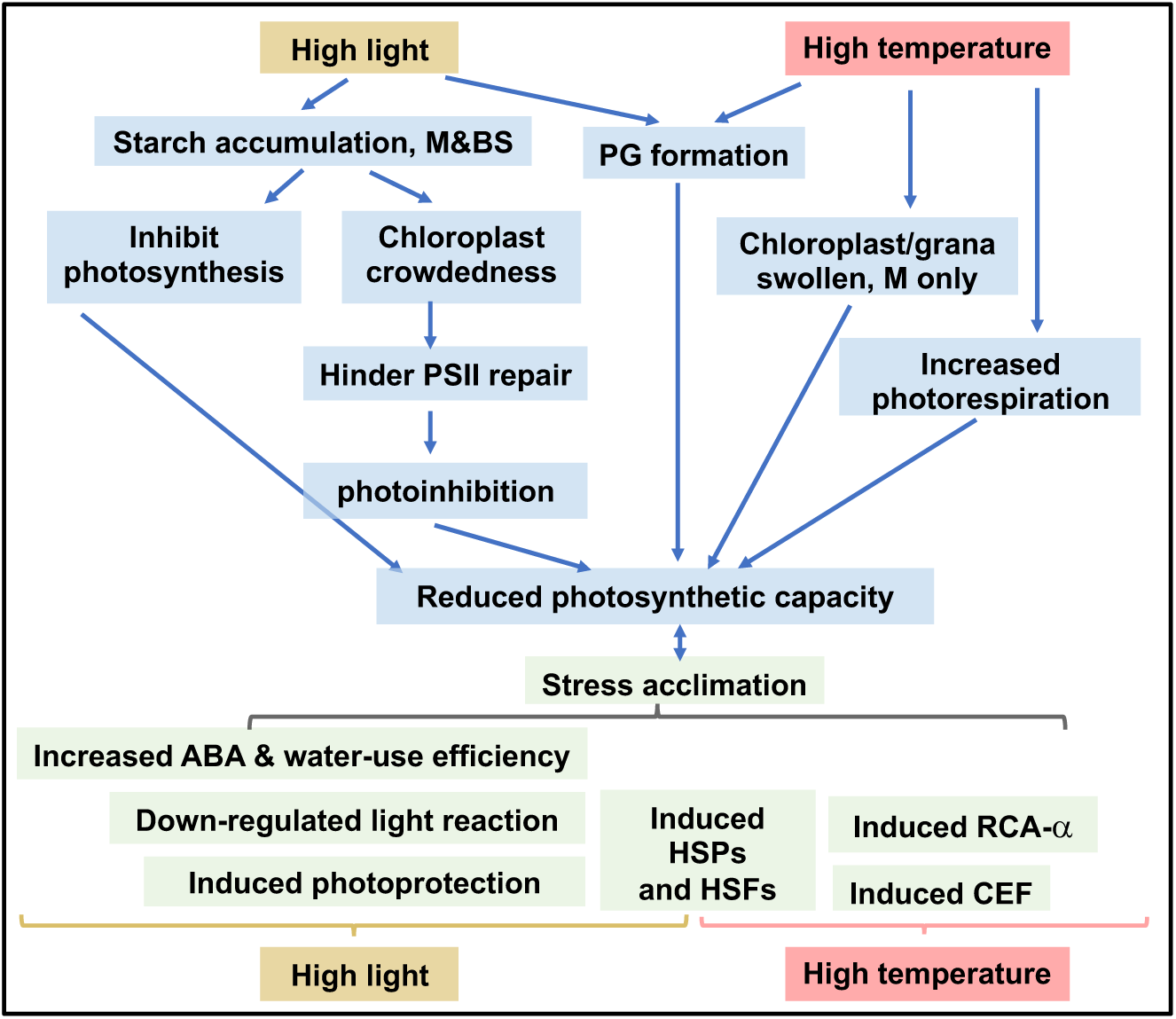
Summary of how *S. viridis* responds to high light (HL) or high temperature (HT). Light blue boxes denote changes that may lead to the reduced photosynthetic capacities; light green boxes denote changes that may be adaptive for stress acclimation. M: mesophyll chloroplasts; BS, bundle sheath chloroplasts. HL-treated leaves had over-accumulated starch and increased chloroplast crowdedness, which may hinder PSII repair and result in photoinhibition. Starch accumulation may also inhibit photosynthesis through feedback regulation. Increased plastoglobuli (PG) formation in HL-treated leaves may affect thylakoid composition and function. Under HT, M chloroplasts had swollen chloroplasts/grana and seem more heat-sensitive than BS chloroplasts. Heat-induced photorespiration and PG formation could further reduce photosynthesis. Meanwhile, HL and HT also induce adaptive responses for acclimation. Under HL, the induced photoprotection, down-regulated light reaction, and increased water-use efficiency through abscisic acid (ABA) can help *S. viridis* acclimate to excess light. Under HT, the induced cyclic electron flow (CEF) and Rubisco activase (RCA-α) can protect photosynthesis from heat stress. The induced heat shock transcription factors (HSFs) and heat shock proteins (HSPs) are adaptive responses to both HL and HT although the induction was much quicker under HT.

### Starch over-accumulation may contribute to photoinhibition in HL-treated leaves

In response to HL, *S. viridis* induced NPQ to dissipate excess light energy via increased *PsbS* transcription and zeaxanthin accumulation (Fig. 3a, 6c). At the transcriptional level, HL-treated plants up-regulated transcripts involved in PSII assembly/repair and photoprotection before down-regulating transcripts involved in LHCII, PSII core complex, and PSI complex (Fig. 3), suggesting a strategy to dissipate light and repair damaged PSII before the remodeling of photosystems. With the rapid induction of photoprotective pathways, it was initially surprising to see the significant amount of photoinhibition in HL-treated leaves of *S. viridis* (Fig. 1b), but the HL-induced starch accumulation may provide some insight.

Our TEM data showed that the mean relative starch volume fraction was increased significantly in both M and BS chloroplasts in HL_4h leaves as compared to ctrl_4h leaves (Fig. 7, Supplementary Fig. 15). The increased starch accumulation likely resulted from increased CO_2_ fixation rates (Fig. 8a) but imbalance of starch synthesis/ degradation and sugar transport from downstream pathways under HL. In C_3_ plants, starch is mostly present in M chloroplasts where photosynthesis occurs^66,67^. In C_4_ plants, starch is present in both BS and M chloroplasts (Fig. 7a-f), although Rubisco dominantly localizes in the BS chloroplasts^67^. The over-accumulated starch increased the crowdedness of the chloroplasts (Fig. 7, Supplementary Fig. 15), which may hinder PSII repair, especially in M chloroplasts where PSII is enriched. PSII complexes are concentrated in the stacked grana regions; during PSII repair, damaged PSII subunits migrate from the stacked grana region to the grana margin and the unstacked grana region (stroma lamellae) where the proteins involved in PSII repair are localized (e.g., FtsH, Deg proteases that degrade damaged PSII subunits)^15,68^. In Arabidopsis under HL, the grana lumen and margin swell to facilitate protein diffusion and PSII repair^23,69^, however, we did not see these changes in HL-treated *S. viridis* leaves (Supplementary Fig. 12d, e, i). Starch overaccumulation and increased chloroplast crowdedness may slow down the migration of damaged PSII subunits and inhibit PSII repair, contributing to the HL-induced photoinhibition (Fig. 1b, 10). Additionally, ATP synthase activity was significantly reduced in HL_4h leaves as compared to ctrl_4h leaves (Fig. 9b,c), which may be associated with the starch accumulation and sugar feedback inhibition of photosynthesis. HL-treated Arabidopsis plants had reduced starch in chloroplasts^70^, which may reflect the differences in experimental conditions or the stronger capability to use HL for carbon fixation in C_4_ plants than C_3_ plants.

### HL differentially regulated genes involved in sugar-sensing pathways

Sugar signaling integrates sugar production with environmental cues to regulate photosynthesis^35,71,72^. In C_3_ plants, some of the sugar-sensing pathways include: (1) SnRK1 pathway (sucrose-non-fermenting 1 related protein kinase 1, starvation sensor, active under stressful and sugar deprivation conditions to suppress growth and promote survival)^73–75^; (2) Trehalose pathway (trehalose is a signal metabolite in plants under abiotic stresses and helps plants survive stresses)^65,76^. Sugar sensing pathways under abiotic stresses are underexplored in C_4_ plants^35^. Our RNA-seq data showed that two subunits of SnRK1 (β*2,* γ*4*) were highly down-regulated under HL (Fig. 4b), suggesting possible inhibition of the SnRK1 pathway. In the trehalose pathway, trehalose-6-phosphate synthase (TPS) produces trehalose-6-phosphate (T6P); the T6P phosphatase (TPP) dephosphorylates T6P to generate trehalose^65^. A copy of the potential catalytically active *TPS* (*TPSI)* in *S. virids* was induced and two copies of *TTP* were down-regulated during HL (Fig. 4b), suggesting possible increased level of T6P. T6P is a signal of sucrose availability, inhibits SnRK1 pathway, promotes plant growth and development^77,78^. Based on the expression pattern of genes involved in sugar-sensing pathways and the over-accumulated starch under HL, we postulated that HL-treated *S. viridis* leaves had increased sugar levels, and possibly up-regulated T6P sugar-sensing pathway to down-regulate the SnRK1 pathway and promote plant growth^76,79^, which may alleviate the stress of starch over-accumulation and photosynthesis inhibition under HL.

### Potential links between HL response and ABA pathway exist in *S. viridis*

The links between HL responses and ABA have been reported in C_3_ plants^11,12,80,81^. Arabidopsis ABA biosynthesis mutants (e.g., *nced3*) were more sensitive to HL than WT^11,12^. HL-treated *S. viridis* leaves had reduced capacity for stomatal conductance (Fig. 8b), which can most likely be attributed to an acute increase of ABA levels in HL-treated leaves (Fig. 5b). Although ABA levels were only significantly increased at HL_1h and then gradually decreased, the ABA-induced stomatal closure may be prolonged. Consistent with this, RNA-seq data showed increased expression of genes involved in ABA responses and signaling during the 4-h HL treatment (Fig. 5a). Stomatal conductance increases with light to increase CO_2_ uptake, which also increases water loss. To reduce water loss and improve water use efficiency, a relatively lower stomatal conductance under HL may be an adaptive response. Our results in *S. viridis* provide insight into the reduced stomatal conductance and photosynthesis in sorghum leaves under HL^9^.

ABA homoeostasis is maintained by the balance of its biosynthesis, catabolism, reversible glycosylation, and transport pathways^19^. Several ABA biosynthesis genes were up-regulated during HL (Fig. 5a), including *NCED1* (9-cis epoxycarotenoid dioxygenase)^19,82,83^ and *ABA1/ZEP1*, suggesting that local, *de novo* ABA biosynthesis may be one source of the rapid and large induction of ABA at HL_1h. The up-regulation of *CYP707As*, which are responsible for ABA degradation^84^, may contribute to the gradual reduction of ABA levels after 1 h HL. Furthermore, the *S. viridis* homolog of Arabidopsis BG1 (glucosidases, hydrolyze inactive ABA-GE to active ABA in endoplasmic reticulum)^85^ was induced at HL_2h and HL_4h. Dehydration rapidly induces polymerization of AtBG1 and a 4-fold increase in its enzymatic activity^85^. It is possible that the hydrolysis of ABA-GE to ABA by polymerized BG1 may precede the induction of the BG1 transcript, contributing to the transiently increased ABA levels. Several putative ABA transporters were not differentially expressed (Supplementary Data 6), but a *S. viridis* homolog of the Arabidopsis ABA importer *ABCG40* was down-regulated in HL (Fig. 5a), suggesting ABA import from other parts of the plant to leaves may be less likely. Thus, the HL increased ABA level may be due to ABA *de novo* biosynthesis and/or reversible glycosylation from ABA-GE to ABA.

### HT responses had distinct features in comparison to HL

Compared to HL, HT_4h leaves showed much less change in starch accumulation, little change in chloroplast crowdedness (Fig. 7), and no photoinhibition (Fig. 1). Under HT, M chloroplasts had reduced relative starch volume fraction but increased chloroplast area as compared to ctrl (Supplementary Fig. 13), suggesting heat-induced chloroplast swelling that is independent of starch accumulation. Grana dimension increased in HT-treated M chloroplasts (Supplementary Fig. 13), suggesting heat-induced grana swelling. In contrast, BS chloroplasts have slightly increased starch, no change of chloroplast area, but decreased grana dimension under HT, suggesting cell-type specific heat responses. PG formation was highly induced in both M and BS chloroplasts under HT, which may be associated with heat-increased thylakoid membrane leakiness, consistent with previous reports^26,86,87^. Induced chloroplast/grana swelling and PG formation may reflect heat-induced damage to chloroplast ultrastructure, which may contribute to the reduced photosynthetic rates under HT.

The transcriptome changes under HT were less extensive but more dynamic than under HL (Fig. 2-5). HT induced more PG formation than HL (Supplementary Fig. 13), however, surprisingly there were few transcriptional changes of genes encoding proteins that localize to PG under HT (Fig. 4a). These results suggest the heat-induced PG formation may be a direct and physical response of thylakoid membranes to moderately HT and not regulated at the transcriptional level.

Response to HT also showed some unique transcriptional changes that were absent or minimal under HL. First, HT resulted in high and sustained induction of Rubisco activase (*RCA_α*) (Fig. 3c). RCA removes inhibitors from Rubisco, maintains Rubisco activation, and is important for carbon fixation^48,49^. Rubisco is thermostable but RCAs are heat labile, resulting in reduced Rubisco activation and CO_2_ fixation under HT^36^. Plants grown in warm environments usually have RCAs that are more thermotolerant^88–90^. In *S. viridis*, maize, and sorghum, HT induces the protein level of *RCA_α* and the rate of *RCA_α* induction is associated with the recovery rate of Rubisco activation and photosynthesis^91^, suggesting the heat-induced RCA_α may be the thermotolerant isoform. Understanding the function and regulation of RCAs may help improve thermotolerance of photosynthesis in C_4_ plants. Additionally, HT upregulated small *HSPs* much quicker than HL.

Key genes involved in photorespiration (Fig. 3c) and cyclic electron flow (CEF) around PSI (Supplementary Fig. 6a) were up-regulated under HT, suggesting HT-induced photorespiration and CEF. C_4_ plants employ carbon-concentrating mechanisms (CCM) to concentrate CO_2_ around Rubisco and reduce photorespiration in the BS chloroplasts. However, *S. viridis* BS chloroplasts have a small number of grana (Fig. 7d-f), where PSII is present and can be a source of O_2_ production. Photorespiration increases with temperature faster than photosynthesis^30,92^ and HT may also increase the CO_2_ leakiness of BS chloroplasts^38,39^, promoting photorespiration and reducing photosynthesis. CEF generates only ATP without NADPH, balances ATP/NADPH ratio, generates transthylakoid proton motive force (*pmf*), and protects both PSI and PSII from photo-oxidative damage in C_3_ plants^93,94^. Increased CEF activity has been frequently reported under stressful conditions in C_3_ plants^26,95,96^, indicating its important role in stress protection. To compensate for the extra ATP needed for CCM, C_4_ plants are proposed to have high CEF in BS chloroplasts^3,97^. CEF is reported to increase in *S. viridis* under salt stress^98^. The heat-induced CEF could protect photosynthesis under HT by maintaining transthylakoid *pmf* and generating extra ATP.

Frey et al. identified 39 heat-tolerance genes in maize that were significantly associated with heat-tolerance and up-regulated in most of the 8 maize inbred lines^41^. Five *S. viridis* homologs of the maize heat-tolerance genes were also up-regulated in our RNA-seq data under HT, providing potential engineering targets to improve heat tolerance in C_4_ plants (Supplementary Data 5).

Although HL and HT responses had their own unique features, their transcriptional responses had significant overlaps (Fig. 2b). We identified 42 highly induced genes (FC ≥ 5) and 13 highly repressed genes (FC ≤ −5) in both conditions (Supplementary Fig. 5, Supplementary Data 5). The 42 highly induced genes provide potential targets for improving resistance to HL and HT in C_4_ crops, including several putative transcription factors, HSP20/70/90 family proteins, β-amylase, and a putative aquaporin transporter for promoting CO_2_ conductivity in C_4_ plants^3,99,100^. Additionally, *HSFA6B* was induced (2≤ FC ≤5) at both HL and HT. It is reported that HSFA6B operates as a downstream regulator of the ABA-mediated stress response and is involved in thermotolerance in Arabidopsis, wheat, and barley^101,102^. This gene may be involved in regulation of genes that are common to both the HL and HT responses and it would be interesting for further study to generate HL and HT tolerant C_4_ crops.

In comparison to the C_3_ model plant Arabidopsis, the C_4_ model plant *S. viridis* has shared and unique responses under HL and HT. The shared responses include induced NPQ, *PsbS* transcription, zeaxanthin accumulation, PG formation, and ABA levels under HL, and the induced PG formation as well as swollen M chloroplasts under HT. The unique responses in *S. viridis* to HL include the over-accumulated starch in both M and BS chloroplasts and increased chloroplast crowdedness. In HT, the unique responses in *S. viridis* include dynamic transcriptome regulation and different heat sensitivities of M and BS chloroplasts. The reduced photosynthetic capacity under HL or HT also demonstrated the need to increase the tolerance to these two stresses in C_4_ plants.

The different responses in M and BS chloroplasts in *S. viridis* are particularly interesting and warrant further study. We sorted HL or HT induced DEGs into M and BS specific pathways based on previously published M/BS transcriptomes^58^ (Supplementary Fig. 8). Although we cannot rule out some transcripts may have altered cell type specificity under stressful conditions, due to the function specificity of the M and BS cells, a significant fraction of the M and BS specific transcripts likely keep similar cell type specificity under our HL and HT conditions as compared to the published control condition. Our analysis revealed M- and BS-specific transcriptional regulation in response to HL or HT in *S. viridis* (Supplementary Fig. 8). Under HL, the majority of M-specific DEGs related to ROS-scavenging and HSPs were up-regulated while the majority of BS-specific DEGs related to these two pathways were down-regulated, suggesting M cells may require more ROS scavenging and HSPs than BS cells in response to HL, probably due to more ROS production and higher need for maintaining protein homeostasis in M cells than BS cells under HL. In contrast, under HT, many ROS-scavenging DEGs were up-regulated in BS cells but down-regulated in M cells (possibly due to heat-induced photorespiration) while DEGs related to HSPs were up-regulated in both cell types. It is intriguing that HL up-regulated M-specific sugar transporters but down-regulated BS-specific sugar transporters. In Arabidopsis, SWEET16/17 plays a key role in facilitating bidirectional sugar transport along sugar gradient across the tonoplast of vacuoles^103,104^. The homologous copy of SWEET16/17 in *S. viridis* is M-cell specific and was up-regulated in HL (Supplementary Fig. 6b), suggesting SvSWEET16/17 may mediate sugar uptake into vacuoles in response to a high centration of cytosolic sugar level in M cells. The down-regulation of BS-specific SWEETs under HL may indicate feedback inhibition of sugar phloem loading due to unmatched sugar usage in downstream processes^105^.

In summary, we elucidated how the C_4_ model plant *S. viridis* responds to moderately HL or HT at the photosynthetic, transcriptomic, and ultrastructural levels (Supplementary Fig. 14). Our research furthers understanding of how C_4_ plants respond to HL and HT by linking the data from multiple levels, reveals different acclimation strategies to these two stresses in C_4_ plants, discovers unique HL/HT responses in C_4_ plants in comparison to C_3_, demonstrates M/BS cell type specificity under HL or HT, distinguishes adaptive from maladaptive responses, and identifies potential targets to improve abiotic stress tolerance in C_4_ crops.

## Methods

### Plant growth conditions and treatments

*S. viridis* ME034 (also known as ME034v) plants were grown in a controlled environmental chamber under constant 31°C, 50% humidity, ambient CO_2_ conditions, 12 h photoperiod, and leaf level light intensity of 250 μmol photons m^−2^ s^−1^. Similar level of growth light has been used in literatures for *S. viridis* under control conditions^58,98,106^. Seeds were germinated on Jolly Gardener C/V Growing Mix (BGF Supply Company, Oldcastle, OCL50050041) and fertilized with Jack’s 15-5-15 (BGF Supply Company, J.R. Peters Inc., JRP77940) with an Electrical Conductivity (EC) of 1.4. At seven days after sowing (DAS), seedlings were transplanted to 3.14” x 3.18” x 3.27” pots. At 13-DAS, 4 h after light was on in the growth chamber, plants with fourth fully expanded true leaves were selected for 4 h HL (leaf level light intensity of 900 μmol photons m^−2^ s^−1^ and chamber temperature of 31°C) or 4 h HT (chamber temperature of 40°C and leaf level light intensity of 250 μmol photons m^−2^ s^−1^) treatments carried out in separate controlled environmental chambers under 50% humidity and ambient CO_2_ conditions. A separate set of plants remained in the control chamber set to growth conditions. Leaf temperature was stable at 31°C under control and HL treatments while it increased gradually from 31°C to 37°C by the end of 4 h treatment of 40°C (Supplementary Fig. 1).

### Gas-exchange and chlorophyll fluorescence measurements

Leaf-level gas exchange and pulsed amplitude modulated (PAM) chlorophyll *a* fluorescence was measured using a portable gas-exchange system LI-6800 coupled with a Fluorometer head 6800-01 A (LI-COR Biosciences, Lincoln, NE). Fourth fully expanded true leaves of *S.viridis* plants from different treatments were first dark-adapted for 20 min in the LI-6800 chamber to measure maximum PSII efficiency (F_v_/F_m_) under constant CO_2_ partial pressure of 400 ppm in the sample cell, leaf temperature 25°C, leaf VPD 1.5 kPa, fan speed 10,000 RPM, and flow rate 500 μmol s^−1^. We then performed the light response curves followed by CO_2_ response curves (*A/C_i_* curve) as described (Supplementary Fig. 2). Red-blue actinic light (90%/10%) and 3-6 biological replicates for each treatment were used for all measurements. We used leaf temperature of 25°C for light and CO_2_ response curves as described in previous publications for *S. viridis* regardless of growth temperatures^56,98, 107–109^. During all measurements, the instrument parameters were consistent and stable. For CO_2_ response curves, all net CO_2_ assimilation rates were corrected with the empty chamber data to count for inevitable and minor LI-6800 leaf chamber leakiness during the CO_2_ response curves following the established protocols^110^.

Photosynthetic parameters were calculated as described^62^ (see formulas, Supplementary Table 1). To estimate the true NPQ, F_m_ used in the NPQ formula (F_m_ / F_m_’ – 1) needs to be the maximum chlorophyll fluorescence in fully relaxed, dark-adapted leaves in which there is no quenching^62,111^. F_m_ and F_m_’ are the maximum chlorophyll fluorescence yields in dark-adapted and light-adapted leaves, respectively^62,111,112^. In ctrl leaves, F_m_ could be reached with 20 min dark-adaptation without further change after that, but HL-treated leaves needed a much longer recovery period to relax the quenching processes due to the light-induced photoinhibition (Supplementary Fig. 9a). Because the values of F_m_ in dark-adapted ctrl_4h leaves were highly consistent among different biological replicates and reflected the reference level of F_m_ (*i.e.*, without stress treatments), we used the mean F_m_ of ctrl_4h leaves as a baseline to calculate NPQ in leaves with different treatments.

To investigate photosynthetic performance in plants immediately following 4 h of different treatments (ctrl, HL or HT), we also performed short LI-6800 measurements for 5 min on each plant immediately after 4 h treatments without dark-adaptation at 400 ppm CO_2_ with indicated leaf temperatures and light intensities (Fig. 8). To estimate photosynthetic parameters under different treatments as in the growth chambers, the LI-6800 leaf chamber was set to simulate the condition of different treatments: ctrl (31°C, 200 µmol photons m^−2^ s^−1^ light), HL (31°C, 600 µmol photons m^−2^ s^−1^ light,) or HT (40°C, 200 µmol photons m^−2^ s^−1^ light). The temperature and light refer to the conditions in the LI-6800 leaf chamber. The light in LI-6800 leaf chamber (90% red and 10% blue) was different from the white light in the growth chamber, therefore we selected two lights in the LI-6800 leaf chamber that were close to the white lights in growth chambers based on the light quantification in the red (580-670 nm) and blue (440-540 nm) spectrum range. LI-6800 light intensities of 200 and 600 µmol photons m^−2^ s^−1^ were also two of the conditions used in the light response curves with dark-adaptation (Fig. 1c, 8, group 1), allowing for direct comparison. Individual plants were used for each measurement and replicate.

The high abundance of PSI in BS chloroplasts of C_4_ leaves can affect chlorophyll fluorescence measurement (up to 50%) and underestimate the PSII efficiency (F_v_/F_m_) and electron transport rates^113,114^. Thus, our chlorophyll fluorescence data were corrected with 0.5 F_o_, which is the mean minimal chlorophyll fluorescence in dark-adapted leaves under the control condition (ctrl_4h). The PSII operating efficiency calculated from the corrected and uncorrected chlorophyll fluorescence data correlated with each other but the corrected data yielded higher PSII efficiency, with the maximum PSII efficiency in ctrl_4h leaves closer to the theoretical values of 0.86^115^ (Fig. 1b).

### Modeling of photosynthetic parameters using leaf-level gas exchange information

To model photosynthetic parameters, we used gas exchange data from light response curves and CO_2_ response curves (*A/c_i_* curves). The model parameterization and analyses were conducted in R 3.4.3 Project software® (R Development Core Team 2016). First, light response curves were fitted as previously described^116^. We fit a non-linear least squares regression (non-rectangular hyperbola) to estimate photosynthetic parameters (Supplementary Fig. 4). *A/c_i_* curves were fitted as previously described^117^ to estimate the *V_cmax_* (the maximum rate of carboxylation). Feng et al. (2013) followed the C_4_ photosynthesis model using a Bayesian analysis approach^118^. The normality of the data was verified with the Shapiro-Wilk test. Statistical analysis was performed using Student’s two-tailed t-test with unequal variance by comparing ctrl_4h with all other conditions.

### RNA isolation

To isolate RNA from leaves, four biological replicates containing two 2-cm mid-leaf segments from two plants for each time point and treatment were collected from fourth fully expanded true leaves into screw cap tubes (USA Scientific, 1420-9700) with a grinding bead (Advanced Materials, 4039GM-S050) and immediately frozen in liquid nitrogen and stored at −80°C. Frozen samples were homogenized using a paint shaker. RNA was extracted using a Trizol method with all centrifugation at 4°C and 11,000 RCF. One mL of Trizol Reagent (Invitrogen, 15596018) was added to homogenized leaf tissue and resuspended, then 200 µL of Chloroform:Isoamyl alcohol (25:1) was added and vortexed. Tubes were centrifuged for 15 min. 600 µL from the aqueous layer was transferred to a clean tube with equal volume Chloroform:Isoamyl alcohol, vortexed, and centrifuged for 5 min. Next, 450 µL of aqueous layer was transferred to 0.7x volume 100% Isopropanol, mixed well, and chilled for 30 min in −20°C freezer. Samples were centrifuged for 15 min to pellet RNA. Supernatant was decanted, and RNA pellet was rinsed twice with ice-cold 75% ethanol with a 2-min centrifugation following each rinse. RNA was dried in a laminar flow hood until residual ethanol evaporated and was resuspended in 50 µL of nuclease free H_2_O. RNA was quantified using a NanoDrop and Qubit RNA Broad Range (BR) Assay Kit (Thermo Fisher Scientific Inc., Q10210) with the Qubit 3.0 machine. RNA integrity was verified using a Bioanalyzer Nano Assay (Genome Technology Access Center, Washington University in St. Louis).

### RNA-seq library construction and sequencing

RNA samples were diluted to 200 ng/µL in nuclease free H_2_O for a total of 1 µg RNA. Libraries were generated with the Quantseq 3’ mRNA-seq library prep kit FWD for Illumina (Lexogen, 015.96). Libraries were generated according to manufacturer’s instructions. Cycle count for library amplification for 1 µg mRNA was tested using the PCR add-on kit for Illumina (Lexogen, 020.96). qPCR was performed and a cycle count of 13 was determined for the amplification of all libraries. For library amplification, the Lexogen i5 6 nt Dual Indexing Add-on Kit (5001-5004) (Lexogen, 047.4×96) was used in addition to the standard kit to allow all libraries to have a unique combination of i5 and i7 indices. All libraries were quantified using Qubit dsDNA High Sensitivity (HS) Assay Kit (Thermo Fisher Scientific Inc., Q32854) with the Qubit 3.0 machine. Prepared libraries were pooled to equimolar concentrations based on Qubit assay reads. Pooled libraries were submitted to Novogene to be sequenced on the HiSeq4000 platform (Illumina) with paired end, 150 bp reads.

### Mapping and transcript quantification

Single-end reads were trimmed and quality-checked using Trim Galore (version 0.6.2). Trimmed reads from each library were mapped and processed for transcript quantification using Salmon (version 1.1.0) in quasi-mapping mode with a transcriptome index built from the *S. viridis* transcript and genome files (Sviridis_311_v2; Phytozome v12.1)^42^. Salmon outputs were imported into R using the Bioconductor package tximport (1.16.0) to extract gene-level expression values represented by transcript per million (TPM) for each gene across every time point, tissue, and treatment group sampled. Principal component analysis was performed with TPM normalized read counts of all genes using the R package FactoMineR^119^.

### Differential expression analysis

Genes that met minimum read count cutoffs of at least 10 raw reads in at least 10% of samples (14,302 genes) were included in differential expression analysis using DeSeq2, FDR < 0.05^120^. HL or HT treatment time points were compared to the control condition from the same time point. Differentially expressed genes between different time points in either HL or HT were visualized in UpSetR^121^. To identify genes in key pathways of interest in *S. viridis*, we used the MapMan annotations for the closely related *S. italica* (<u>RRID:SCR_003543</u>). From the *S. italica* MapMan annotations, we identified the best hit in *S. viridis* for genes in pathways of interest. We then manually curated these lists based on relevant literature to obtain genes in pathways of interest (Supplementary Data 6), as well as to provide further annotation information for genes identified using the MapMan annotations. We sorted the differentially expressed genes in pathways of interest into fold change (FC) bins based on their DeSeq2 fold change values and presented their expression patterns. FC bins were defined as follows: highly induced: FC ≥ 5; moderately induced: 5 > FC ≥ 2; slightly induced: 2 > FC > 0; not differentially expressed: FC = 0; slightly repressed: 0 > FC > −2; moderately repressed: −2 ≥ FC > −5; highly repressed: FC ≤ −5. Heatmaps of pathways of interest were generated using the R package pheatmap (version 1.0.12. https://CRAN.R-project.org/package=pheatmap).

### ABA quantification

Leaf samples of three biological replicates were harvested at 0, 1, 2 and 4 hours of ctrl, HL, or HT treatment. The fresh leaf weight was immediately measured after harvesting. The samples were quickly placed in liquid nitrogen and then stored in −80°C freezer until further processing. Frozen leaf tissue was homogenized and 15 ng of [^2^H_6_]-abscisic acid was added as an internal standard. Samples were dried to completeness under vacuum. ABA was resuspended in 200µl of 2% acetic acid in water (v/v) and then centrifuged; an aliquot was then taken for quantification. Foliar ABA levels were quantified by liquid chromatography tandem mass spectrometry with an added internal standard using an Agilent 6400 Series Triple Quadrupole liquid chromatograph associated with a tandem mass spectrometer according to the previously described methods^122^.

### Pigment analysis

Three biological replicates of one 2-cm middle leaf segment were collected from fourth fully expanded true leaves into screw cap tubes (USA Scientific, 1420-9700) with a grinding bead (Advanced Materials, 4039GM-S050), immediately frozen in liquid nitrogen, and stored at −80°C. During pigment extraction, 600 µl ice-cold acetone were added to the samples before they were homogenized in a FastPrep-24 5G (MP Biomedicals) at 6.5 m/s for 30 s at room temperature. Cell debris were removed by centrifugation at 21,000 x g for 1 min. The supernatant was filtered through a 4 mm nylon glass syringe prefilter with 0.45 µm pore size (Thermo Scientific) and analyzed by HPLC. HPLC analyses were performed on an Agilent 1100 separation module equipped with a G1315B diode array and a G1231A fluorescence detector; data were collected and analyzed using Agilent LC Open Lab ChemStation software. Pigment extracts were separated on a ProntoSIL 200-5 C30, 5.0 μm, 250 mm by 4.6 mm column equipped with a ProntoSIL 200-5-C30, 5.0 μm, 20 mm by 4.0 mm guard column (Bischoff Analysentechnik) and gradient conditions as previously described^123^. Assuming interconversion of the intermediate antheraxanthin between both zeaxanthin and violaxanthin, the de-epoxidation level can be calculated by (zeaxanthin + 0.5 antheraxanthin) / (violaxanthin + antheraxanthin + zeaxanthin)^124^.

### Transmission electron microscopy (TEM)

*S. viridis* leaves were collected after 4 h of different treatments and prepared for TEM. Four-millimeter biopsy punches were taken from the middle leaf segments of the fourth fully expanded leaves and fixed for 2 h in 2% paraformaldehyde and 2% glutaraldehyde (EM Science, Hatfield, PA, USA) plus 0.1% Tween20 in 0.1 M sodium cacodylate at pH 7.4 at room temp and then at 4°C overnight. Samples were then rinsed 3x in buffer and fixed in 2% osmium tetroxide (EM Science, Hatfield, PA, USA) in ELGA water for 2 h, rinsed 3x in ELGA water and placed in 1% uranyl acetate in ELGA water at 4°C overnight and then at 50°C for 2 h. Next, samples were rinsed 5x in water, dehydrated in a graded acetone series and embedded in Epon-Araldite (Embed 812, EM Science, Hatfield, PA, USA). Embedments were trimmed and mounted in the vise-chuck of a Leica Ultracut UCT ultramicrotome (Leica, Buffalo Grove, IL, USA). Ultrathin sections (∼60 to 70 nm) were cut using a diamond knife (type ultra 35°C; Diatome), mounted on copper grids (FCFT300-CU-50, VWR, Radnor, PA, USA), and counterstained with lead citrate for 8 min^125^. Samples were imaged with a LEO 912 AB Energy Filter Transmission Electron Microscope (Zeiss, Oberkochen, Germany). Micrographs were acquired with iTEM software (ver. 5.2) (Olympus Soft Imaging Solutions GmbH, Germany) with a TRS 2048 x 2048k slow-scan charge-coupled device (CCD) camera (TRÖNDLE Restlichtverstärkersysteme, Germany). Ninety electron micrographs were quantified for each experimental treatment using image analysis (FIJI software, National Institutes of Health) and stereology (Stereology Analyzer version 4.3.3, ADCIS, France). Each TEM image was acquired at 8,000X magnification and 1.37 nm pixel resolution with arrays of up to 5X5 tiles using automated Multiple Image Alignment software module (settings: correlation =1, FFT algorithm, overlap area = linear weighted, movement = emphasize, and equalize). TEM images were analyzed with Stereology Analyzer software version 4.3.3 to quantify relative volume of various cell parameters including stroma, stroma lamellae, starch granules, and grana within individual chloroplasts (Supplementary Fig. 11b). Grid type was set as “point” with a sampling step of 500×500 pixels and pattern size of 15×15 pixels. The percent of relative volume for each parameter was collected after identifying all grid points within one chloroplast and further analyzed in excel. TEM images with a magnification of 8K were used in the Fiji (ImageJ) analysis. The images were scaled to 0.7299 pixel/nm in ImageJ before analyzing the chloroplast area, plastoglobuli area, and grana dimensions. The height of grana margin (positions 1 and 3) and grana core (position 2) were quantified as described previously^23^ (Supplementary Fig. 12d, e). The “polygon selections” tool was used to quantify the chloroplast and plastoglobuli area by outlining the target structure. The individual plastoglobuli (PG) size was measured using ImageJ. All PG in a chloroplast were quantified to get the total PG area per chloroplast. The “straight” tool was used to quantify grana height and width. The grana number and PG number were counted manually. Choosing the correct statistical test to reflect the quantified data is essential in making conclusions. Three different statistical tests were used to find the significance of p-values. The negative binomial test was used for counting data that followed a negative binomial distribution. The Kolmogorov-Smirnov test was used for relative volume data since it is commonly used to find significance between data in a form of ratios. A two-tailed t-test with unequal variance was used for all other data that followed a normal distribution. All three statistical tests compared the treatment conditions to the ctrl conditions of the same cell type. Each treatment had three biological replicates and a total of 90∼120 images of each treatment were analyzed.

### Starch quantification

To isolate starch from leaves, three biological replicates of 2-cm mid-leaf segments were collected from fourth fully expanded true leaves into screw cap tubes (USA Scientific, 1420-9700) with a grinding bead (Advanced Materials, 4039GM-S050) and immediately frozen in liquid nitrogen and stored at −80°C. Frozen samples were homogenized using a paint shaker. For starch quantification, leaves decolorized by 80% ethanol and starch concentration was subsequently measured using a starch assay kit (Megazyme, K-TSTA-100A).

### MultispeQ measurement

A MultispeQ^61^ v2.0 was used to measure chlorophyll fluorescence parameters and electrochromic shift (ECS) in *S. viridis* leaves at the start or after 4 h treatments of ctrl, HL, or HT. ECS results from light-dark-transition induced electric field effects on carotenoid absorbance bands^62,126^ and is a useful tool to monitor proton fluxes and the transthylakoid proton motive force (*pmf*) *in vivo*^63,64^. Light drives photosynthetic electron transport along the thylakoid membrane and proton fluxes across the thylakoid membrane. Protons flux into the thylakoid through H_2_O oxidation at PSII and plastoquinol oxidation at cytochrome *b_6_f* complex; protons flux out of the thylakoid mainly through ATP synthase to make ATP, which is driven by the transthylakoid *pmf*^63,64^. The total amplitude of ECS signal during the light-dark-transition, ECS_t_, represents the transthylakoid *pmf*. The decay time constant of light-dark-transition induced ECS signal, τ_ECS_, is inversely proportional to proton conductivity (ɡ_H_^+^ = 1 / τ_ECS_), which is proportional to the aggregate conductivity (or permeability) of the thylakoid membrane to protons and largely dependent on the activity of ATP synthase^62^. The proton flux rates, v_H+_, calculated by ECS_t /_ τ_ECS_, is the initial decay rate of the ECS signal during the light-dark-transition and reflects the rate of proton translocation by the entire electron transfer chain, usually predominantly through the ATP synthase^62^. ECS was measured using MultispeQ and the dark interval relaxation kinetics with a modified Photosynthesis RIDES protocol at light intensities of 250, 500, and 1000 µmol photons m^−2^ s^−1^. The MultispeQ v2.0 was modified with a light guide mask to improve measurements on smaller leaves. Parameters at the different light intensities were measured sequentially on the middle segment of a fourth fully expanded true leaf at room temperature with no dark adaptation prior to measurements. The estimated NPQ, NPQ_(T),_ was measured by MultispeQ based on a method that does not require a dark-adapted state of the leaf for determination of F_m_^59^. NPQ_(T)_ uses the minimal fluorescence (F_o_’) and maximal fluorescence (F_m_’) in light-adapted leaves to estimate NPQ. Statistical significance was assigned with a two-tailed t-test assuming unequal variance.

### Statistics and reproducibility

All data presented had at least 3 biological replicates. Detailed information about statistics analysis were described for each method above.

### Data availability

The datasets analyzed in this paper are included in this published article and supplementary information files. Other information is available from the corresponding author on request.

## Supporting information

Supplementary_Table_1_photosynthetic_parameters

Supplementary_Data_1_TPM_matrix

Supplementary_Data_2_DeSeq2_output

Supplementary_Data_3_overlapping_genes_within_treatments

Supplementary_Data_4_overlapping_genes_between_treatments

Supplementary_Data_5_highly_induced_or_repressed_genes_HLandHT

Supplementary_Data_6_genes_used_for_heatmaps

Fig7_with_high_resolution_TEM_images

SupFig11_with_high_resolution_TEM_image

## Frequently Used Abbreviations

HL: high light
HT: high temperature
ctrl: control
M: mesophyll
BS: bundle sheath
Rubisco: ribulose-1,5-bisphosphate carboxylase/oxygenase
PSII: photosystem II
PSI: photosystem I
A_Net_: net CO_2_ assimilation rates
DEGs: differentially expressed genes
NPQ: non-photochemical quenching
CEF: cyclic electron flow around PSI
RCA: Rubisco activase
ABA: abscisic acid
PG: plastoglobuli
HSP: heat shock protein
HSF: heat shock transcription factor

## ACKNOWLEDGEMENTS

The research was supported by the Defense Advanced Research Projects Agency (DARPA) (HR001118C0137 to R.Z., A.E., D.N., and R.V.) and start-up funding from Donald Danforth Plant Science Center (DDPSC) to R.Z. E.M. was supported by the William H. Danforth Fellowship in Plant Sciences and Washington University in St. Louis. D.P. was supported by the Berkeley Fellowship and the NSF Graduate Research Fellowship Program Grant DGE 1752814. O.D. was supported by the Deutsche Forschungsgemeinschaft (DFG) - Project number 427925948. K.K.N. is an investigator of the Howard Hughes Medical Institute. We would like to thank Dr. Helmut Kirchhoff and Dr. Charles Pignon for helpful discussion about the TEM data and LI-6800 data, respectively. We also want to thank Drs. Blake Meyers, Sona Pandey, and Ivan Baxter for their valuable feedback about the manuscript. We acknowledge imaging support from the Advanced Bioimaging Laboratory (RRID:SCR_018951) at DDPSC and usage of the LEO 912AB Energy Filter TEM acquired through a National Science Foundation (NSF) Major Research Instrumentation grant (DBI-0116650). We also acknowledge the use of the Metabolite Profiling facility of the Bindley Bioscience Center, a core facility of the National Institute of Health-funded Indiana Clinical and Translational Sciences Institute, for assisting in the quantification of ABA levels. We appreciate the DDPSC Plant Growth Facilities for growth chamber reservation and taking care of our plants. DARPA approved the paper for Public Release, Distribution Unlimited.

## AUTHOR CONTRIBUTIONS

R.Z. supervised the whole project. R.Z. and C.M.A. designed and planned all the experiments. C.M.A. led the project, performed and analyzed all LI-6800 data, extracted RNA and prepared the RNA-seq library, and led sample harvest for pigment analysis. C.M.A. and E.B. grew all plants needed for the project. T.J.A. provided insight to optimize gas exchange and chlorophyll fluorescence measurements in *S. viridis* using the LI-6800. T.T. and R.V. performed modeling of the leaf-level gas exchange data. N.Z., E.B. and K.J.C. performed TEM analysis. N.Z. quantified starch using assay kits. W.M. helped harvest leaf tissues and performed MultispeQ measurements. E.B. harvested leaves for ABA measurements and S.A.M.M. performed ABA analyses. D.P., O.D., and K.K.N. performed leaf pigment analysis by HPLC. M.B. and A.L.E. provided insight for RNA-seq library preparation. J.Y. and A.L.E. preprocessed RNA-seq data. E.M. led RNA-seq data analysis and generated all the heatmaps. S.P. and R.Z. identified ABA-related genes in *S. viridis*. E.M. identified all other genes used for the heatmaps. J.B. provided suggestions for statistical analysis. M.W. and D.A.N. helped plan, coordinate, and discuss the RNA-seq experiments. R.Z., C.M.A., and E.M. led the writing of the manuscript with the contribution of all other co-authors. All the authors discussed the results, contributed to data interpretation, and helped revise the manuscript.

## Supplementary information

**Supplementary Table 1:** Formulas to calculate photosynthetic parameters.

## Supplementary Data Files

**Supplementary Data 1:** Normalized read counts in Transcripts Per Million (TPM) for all genes in all time points and biological replicates. Annotation information includes the *S. viridis* provisional defline, *A. thaliana* and *O. sativa* best hits and deflines from the Joint Genome Institute bulk annotation information.

**Supplementary Data 2:** Differential expression data for all genes that were significantly differentially expressed in at least one time point and condition (DeSeq2, FDR < 0.05). Annotation information includes the *S. viridis* provisional defline, *A. thaliana* and *O. sativa* best hits and deflines from the Joint Genome Institute bulk annotation information.

**Supplementary Data 3:** Genes up-regulated or down-regulated at all time points in high light or high temperature conditions. Annotation information includes the *S. viridis* provisional defline, *A. thaliana* and *O. sativa* best hits and deflines from the Joint Genome Institute bulk annotation information.

**Supplementary Data 4:** Overlapping differentially expressed genes between high light and high temperature conditions. Genes differentially expressed in at least one time point were included in lists of up- and down-regulated genes in each condition. Annotation information includes the *S. viridis* provisional defline, *A. thaliana* and *O. sativa* best hits and deflines from the Joint Genome Institute bulk annotation information.

**Supplementary Data 5:** Genes highly induced or highly repressed (FC ≥ 5, or ≤ −5) in both the high light and high temperature treatments during at least one time point. Annotation information includes the *S. viridis* provisional defline, *A. thaliana* and *O. sativa* best hits and deflines from the Joint Genome Institute bulk annotation information. Additionally, heat tolerance genes identified in maize with homologs in *S. viridis*.

**Supplementary Data 6:** *S. viridis* v2.1 gene information used to generate heatmaps of pathways of interest.

**Supplementary figure 1-14**

---

**Supplementary Figure 1.**
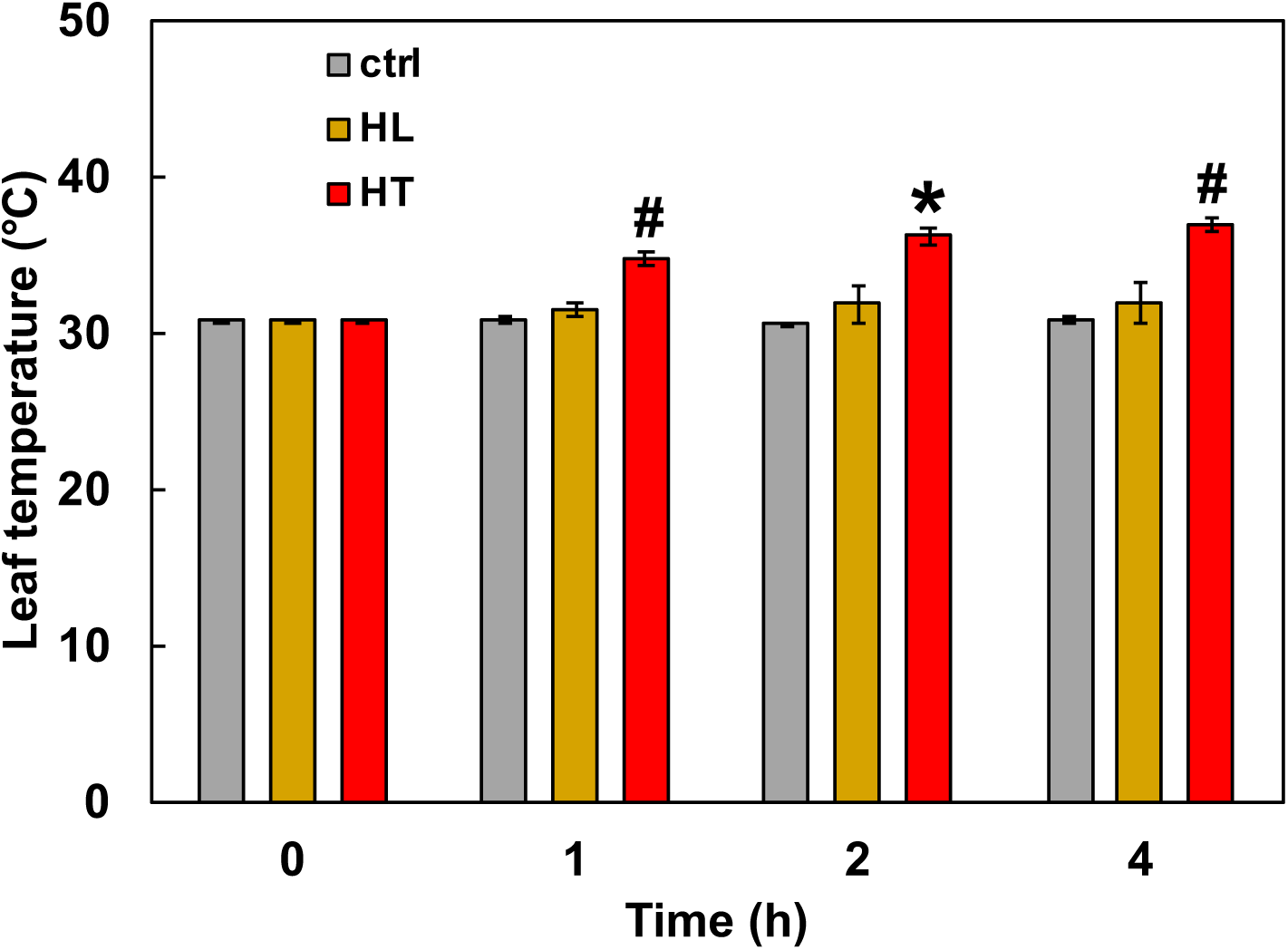
Leaf temperatures of *S. viridis* stayed constant during the control and high light treatments while increased during high temperature treatment. Leaf temperatures of *S. viridis* measured over the 4 h time course of control or high light or high temperature treatments. *S. viridis* plants were treated with control growth condition (ctrl, 31°C and 250 μmol m^−2^ s^−1^ light) or high light (HL, 31°C and 900 μmol m^−2^ s^−1^) or high temperature (HT, 40°C and 250 μmol m^−2^ s^−1^ light) in different growth chambers for 4 h. Leaf temperature was measured at 0, 1, 2, and 4 h for each treatment. Mean ± SE, *n* = 3 biological replicates. Asterisk and pound symbols indicate statistically significant differences of HL and HT compared to ctrl in a given time point using Student’s two-tailed t-test with unequal variance (*0.01<p<0.05, #p<0.01). No significant changes of leaf temperatures during the ctrl and HL condition.

**Supplementary Figure 2.**
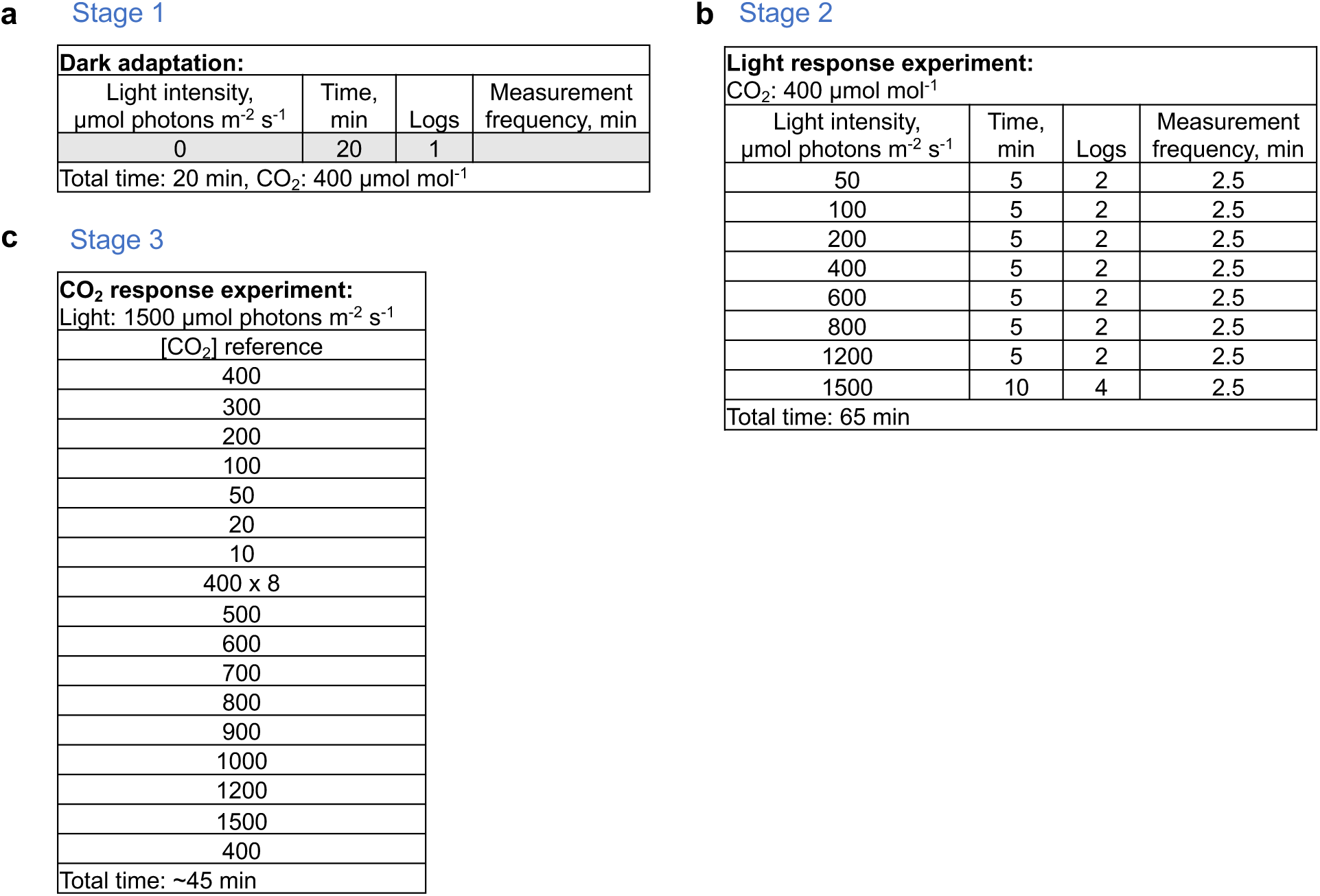
LI-6800 protocol for characterizing photosynthetic parameters. Before or after 4 h different treatments, intact *S. viridis* leaves were dark-adapted in LI-6800 leaf chamber for 20 min (a) to measure the maximum PSII efficiency (F_v_/F_m_), followed by (b) light response experiment from 50 – 1500 μmol photons m^−2^ s^−1^ light and then (c) CO_2_ response experiment at 1500 μmol photons m^−2^ s^−1^ light.

**Supplementary Figure 3:**
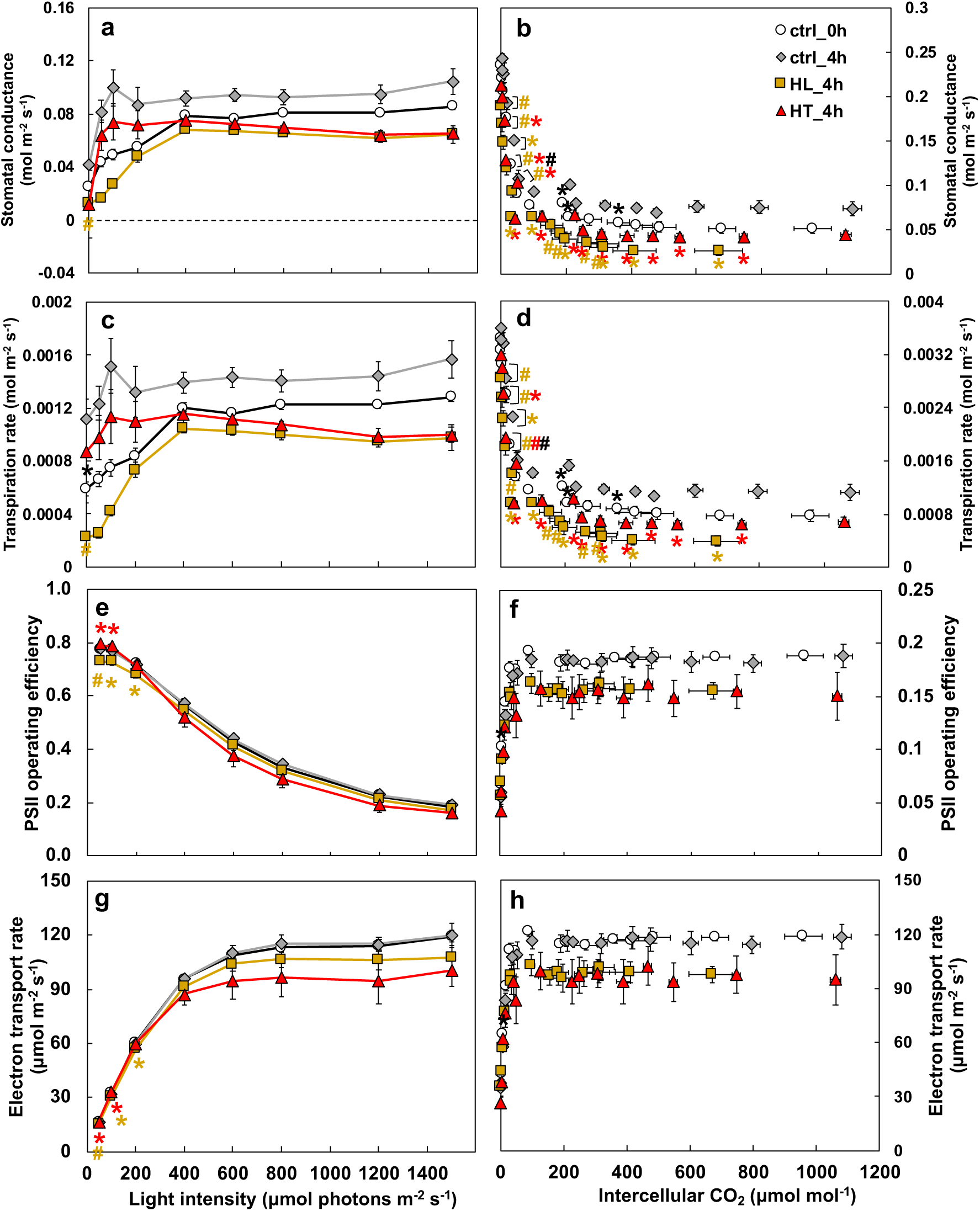
High light or high temperature treatments affected photosynthetic parameters measured by gas exchange and chlorophyll fluorescence. Photosynthetic parameters measured during light (a, c, e, g) and CO_2_ (b, d, f, h) response. Mean ± SE, *n* = 3-6 biological replicates. Asterisk and pound symbols indicate statistically significant differences of ctrl_0h (at the start of treatments), HL_4h (after 4 h high light), and HT_4h (after 4 h temperature) compared to ctrl_4h (after 4 h control treatment) using Student’s two-tailed t-test with unequal variance. P-values were corrected for multiple comparisons using FDR (*0.01<p<0.05, #p< 0.01, the colors of * and # match the significance of the indicated conditions, black for ctrl_0h, yellow for HL_4h, red for HT_4h).

**Supplementary Figure 4:**
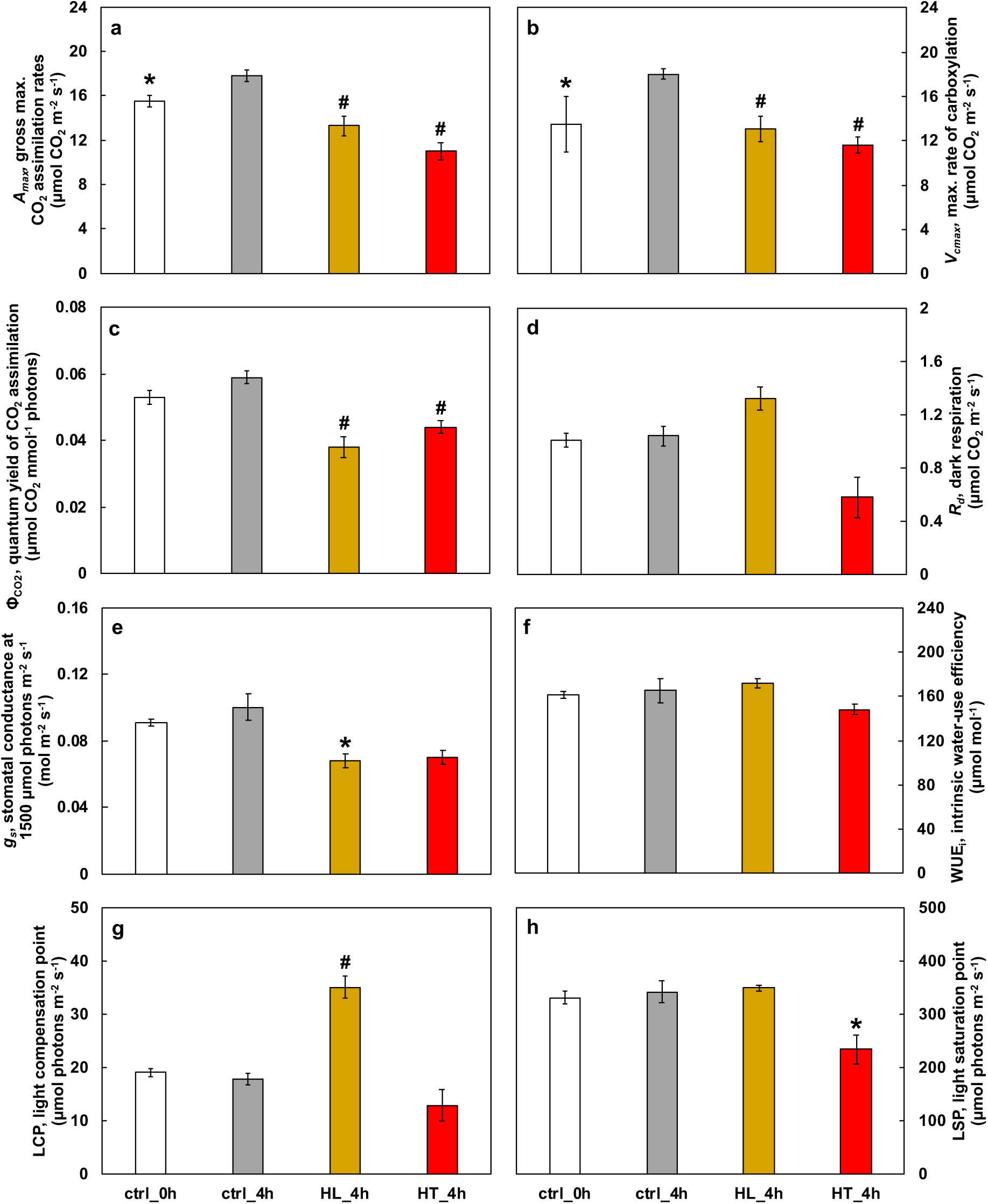
High light or high temperature treated leaves had reduced photosynthetic efficiency. Photosynthetic parameters were derived from light and CO_2_ response curves (mean ± SE; *n* = 3-6). (a) the maximum gross CO_2_ assimilation rates, *A_max_*; (b) the maximum rate of carboxylation, *V_cmax_*; (c) the quantum yield of CO_2_ assimilation, Φ_CO2_, which is the ratio of the moles of CO_2_ fixed in photosynthesis per mole of quanta (photons of light) absorbed, and is a measure of the efficiency in which light is converted into fixed carbon; (d) the day-time dark respiration rate, *R_d_*, equal to *A_n_* when light intensity is zero; (e) stomatal conductance, *g_s_*; (f) water use efficiency, WUE; (g) light compensation point, LCP, the threshold of low light intensity at which photosynthesis is equal to leaf respiration and, therefore *A_n_* is zero; (h) light saturation point, LSP, the estimated light intensity where 75 % of *A_max_* was reached. Asterisk and pound symbols indicate statistically significant differences of ctrl_0h (at the start of treatments), HL_4h (after 4 h high light), and HT_4h (after 4 h temperature) compared to ctrl_4h (after 4 h control treatment) using Student’s two-tailed t-test with unequal variance (*0.01<p<0.05, #p< 0.01).

**Supplementary Figure 5.**
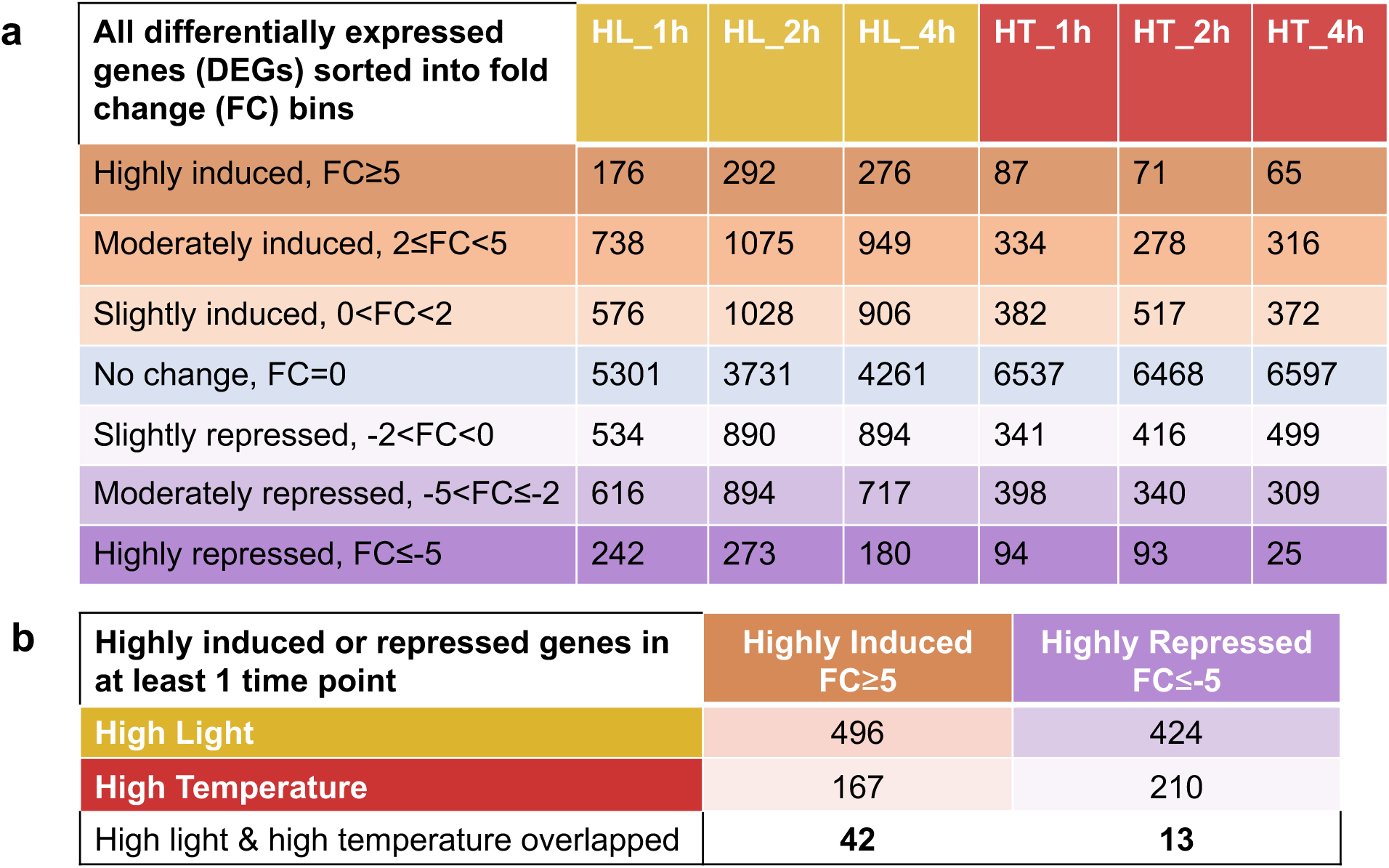
High light had more differentially regulated genes (DEGs) than high temperature while both also had overlapping DEGs. **(a)** Table of DEGs sorted into bins based on their DeSeq2 fold change values at each time point of high light (HL) or high temperature (HT) treatment. All genes that are differentially expressed in at least one timepoint are shown, those that are not differentially expressed at a given time point are represented in the “No change, FC = 0” category. **(b)** Number of genes that are highly induced (FC ≥ 5) or highly repressed (FC ≤ −5) in at least one time point in either HL or HT treatment. 42 genes are highly induced in at least one timepoint in both HL and HT and 13 genes are highly repressed in both HL and HT.

**Supplementary Figure 6.**
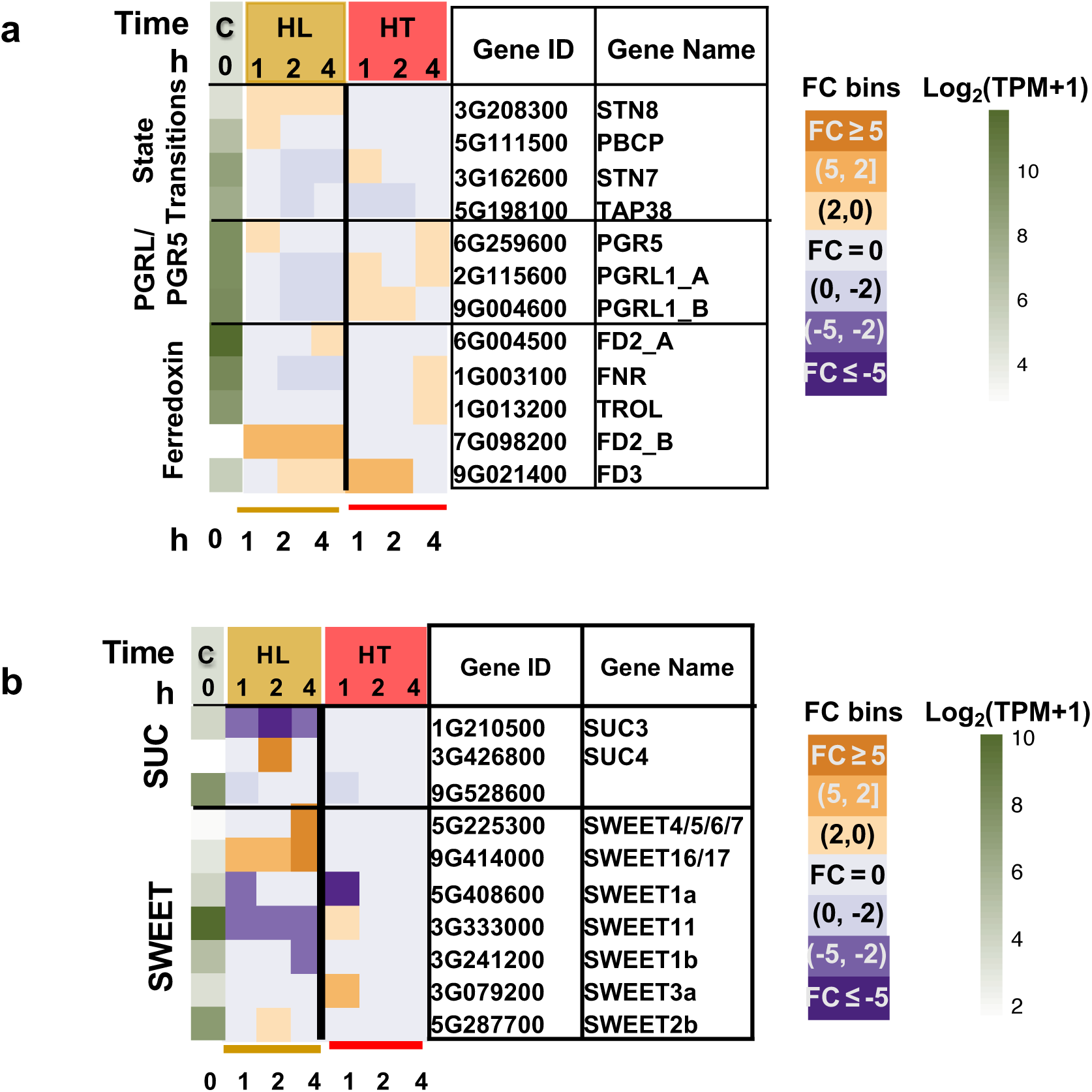
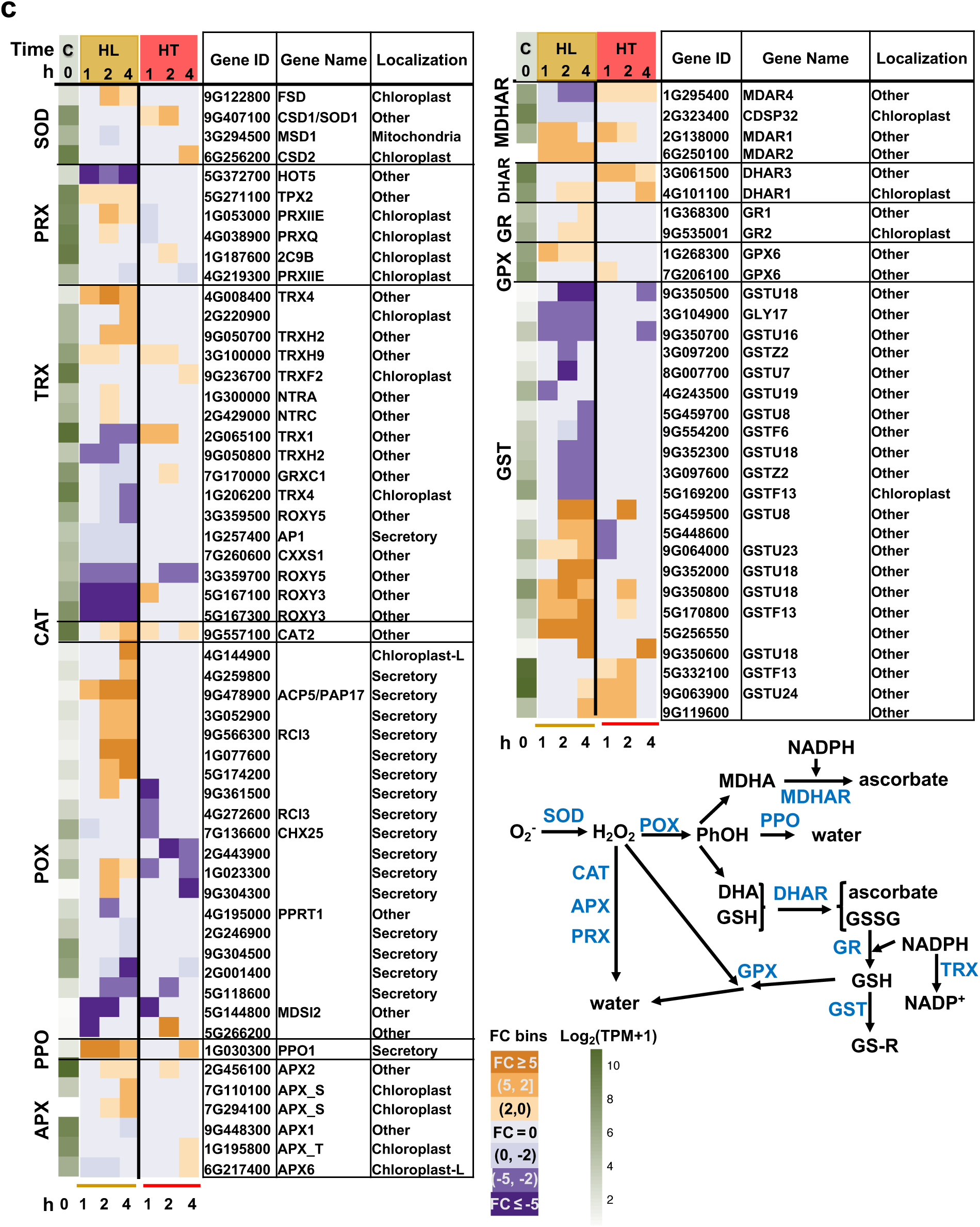
High light (HL) or high temperature (HT) differentially regulated genes involved in various pathways associated with photosynthesis. **(a)** Alternative light reactions of photosynthesis. *PGR5 (proton gradient regulation 5)* and *PGRL1 (PGR5-like photosynthetic phenotype 1)* are genes involved in cyclic electron transport around photosystem I. **(b)** Genes encoding sugar transporters. *SUC (Sucrose-proton symporters)* and *SWEET (Sugar Will Eventually Exported Transporters)* encode sucrose transporters. **(c)** Genes involved in antioxidant defense pathways. SOD: superoxide dismutase; PRX: peroxiredoxins; TRX: thioredoxin; CAT: catalase. POX: peroxidases. PPO: polyphenol oxidase; APX: ascorbate peroxidase; MDHAR: monodehydroascorbate reductase; DHAR: dehydroascorbate reductase; GR: glutathione reductase; GPX: glutathione peroxidase; GST: glutathione S-transferase. These antioxidant enzymes are colored in blue in the antioxidant defense pathways based on Hasanuzzaman et al^57^. SOD leads the frontline defense in the antioxidant defense system by converting superoxide anion (O_2_) into hydrogen peroxide (H_2_O_2_) which is further detoxified to water (H_2_O) by one of these enzymes: POX, CAT, APX, PRX, or GPX. MDHA, monodehydroascorbate; PhOH, phenolic compounds; DHA, dehydroascorbate; GSH, reduced Glutathione; GSSG, oxidized glutathione; R, aliphatic, aromatic, or heterocyclic group; NADPH, nicotinamide adenine dinucleotide phosphate. Most antioxidant enzymes have multiple gene family members in *S. viridis*. The first green column displays log_2_(mean TPM + 1) at ctrl_0h (at the start of treatments, C). TPM, transcripts per million, normalized read counts. Heatmap displays the fold change (FC) bin of DeSeq2 model output values at 1, 2, 4 h of HL or HT versus control at the same timepoint (q < 0.05). FC bins: highly induced: FC ≥ 5; moderately induced: 5 > FC ≥ 2; slightly induced: 2 > FC > 0; not differentially expressed: FC = 0; slightly repressed: 0 > FC > −2; moderately repressed: −2 ≥ FC > −5; highly repressed: FC ≤ −5. Gene ID: *S. viridis* v2.1 gene ID, excluding “Sevir.”. All genes presented in the heatmaps were significantly differentially regulated in at least one time point.

**Supplementary Figure 7.**
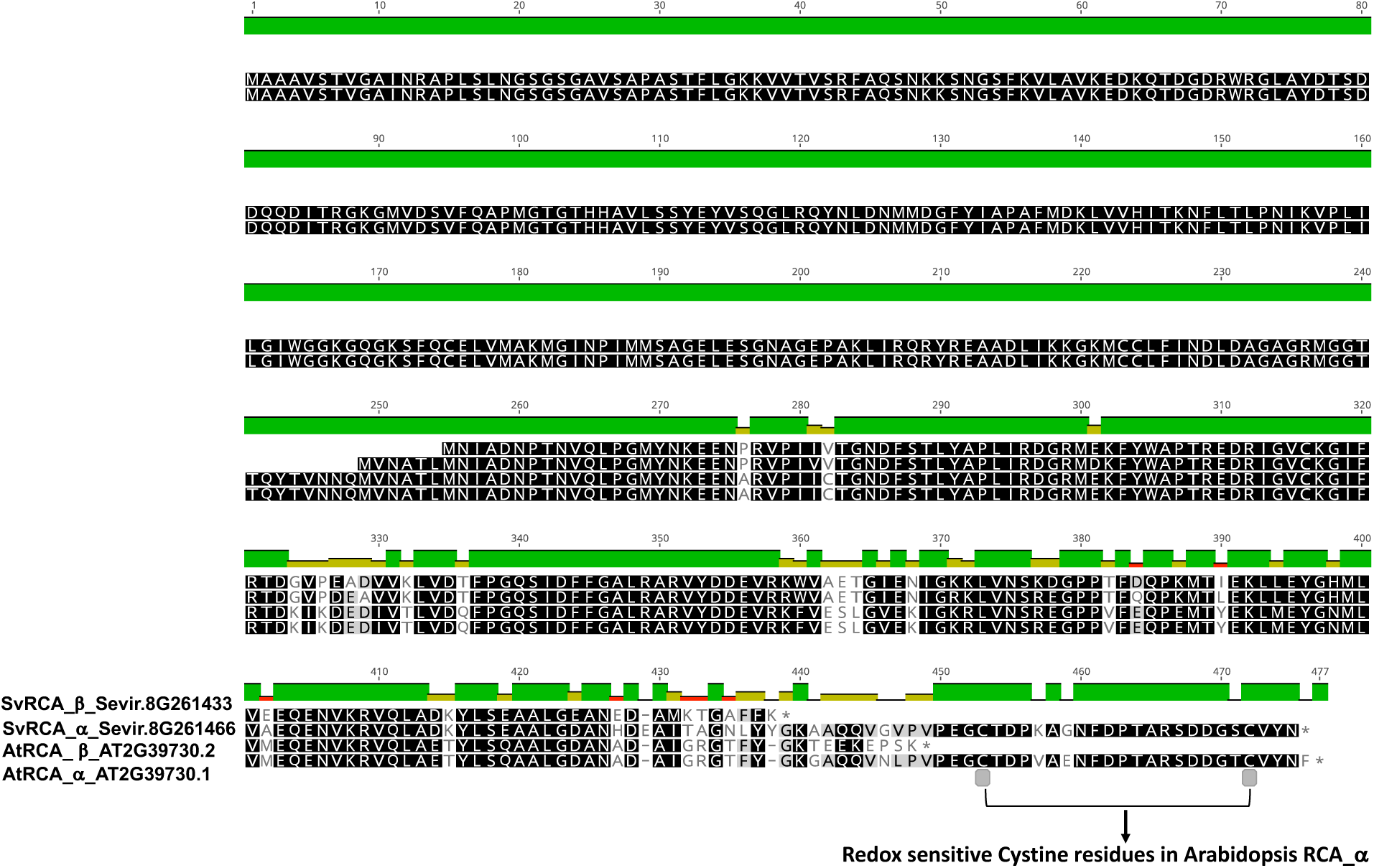
Peptide sequence alignment of two *S. viridis* Rubisco Activases (RCAs) with *A. thaliana* RCAs reveals α and β copies of RCA in *S. viridis*. *A. thaliana* RCA_α has two redox sensitive cysteine residues, which are retained in the *S. viridis* RCA_α copy.

**Supplementary Figure 8.**
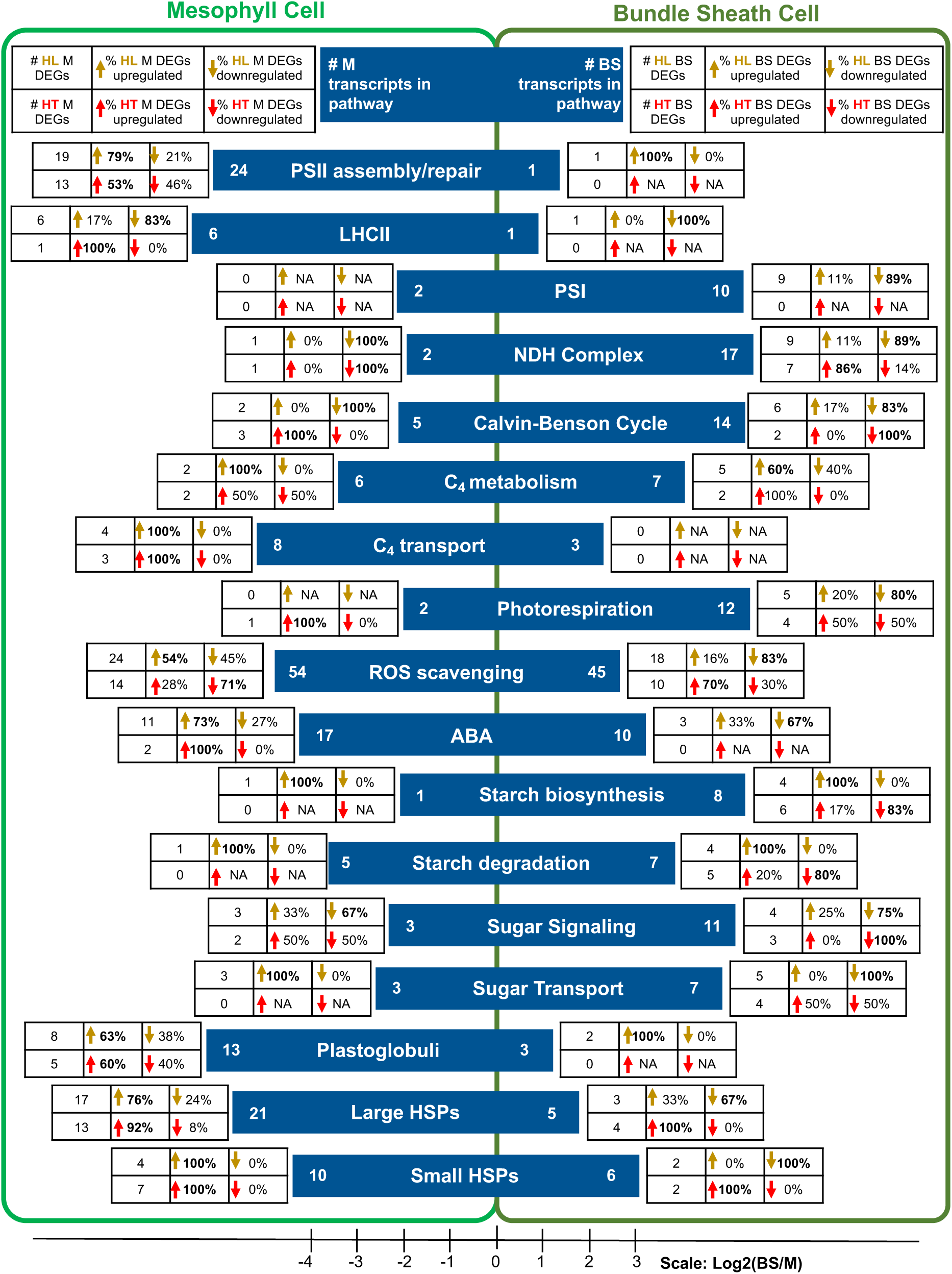
Mesophyll (M) and bundle sheath (BS) specificity of differentially expressed genes reveals cell type specific responses to high light (HL) or high temperature (HT). Light and dark green box denotes M and BS cells, respectively. The blue horizontal bars denote pathways of interest we investigated. Position of blue horizontal bars indicates cell type specificity of pathways, and represents the Log2(number of BS specific transcripts/number of M specific transcripts associated with a pathway) according to the published M and BS specific transcriptome data in *S. viridis* under the control condition^58^. The numbers at the left and right end of each blue horizontal bar represent the numbers of M or BS specific transcripts associated with this pathway. Pathways that are preferentially expressed in M cells (more M than BS specific transcripts, log2(BS/M)<0) under control conditions include PSII assembly, LHCII, C_4_ transport, ROS scavenging, ABA, PG, and HSPs. Pathways that are preferentially expressed in BS cells (more BS than M specific transcripts, log2(BS/M)>0) under control conditions include PSI, NDH complex, Calvin-Benson cycle, C_4_ metabolism, photorespiration, starch biosynthesis/degradation, sugar signaling, and sugar transport. Legends for M and BS specific transcript data in responses to HL or HT are on the top part of the light and dark green box, respectively. Each pathway has a table of data for each cell type. For each table, the first column indicates the number of M or BS specific transcripts related to a pathway that were differentially expressed in HL (top) or HT (bottom) in at least one time point. The rest two columns of the table represent the fraction of up- (2^nd^ Col) or down-regulated (3^rd^ Col) transcripts out of the total number of cell-type specific differentially expressed genes (DEGs) related to a pathway in HL (top) or HT (bottom). Bolded percentage indicates the larger portion (either up- or down-regulated) in HL or HT in each cell type. In HL, 83% of the BS-specific ROS-scavenging DEGs were down-regulated, whereas 54% of M-specific ROS-scavenging DEGs were up-regulated. In contrast, in HT, the majority of BS-specific ROS-scavenging DEGs were up-regulated while M-specific ROS-scavenging DEGs were down-regulated, which may be related to heat-induced photorespiration in BS chloroplasts. In HL, all differentially expressed sugar transports were up-regulated in M cells but down-regulated in BS cells. For HSPs, the majority of M-specific DEGs were up-regulated while the majority of BS-specific DEGs were down-regulated in HL. However, in HT, the majority DEGs of HSPs in both M and BS cells were upregulated.

**Supplementary Figure 9.**
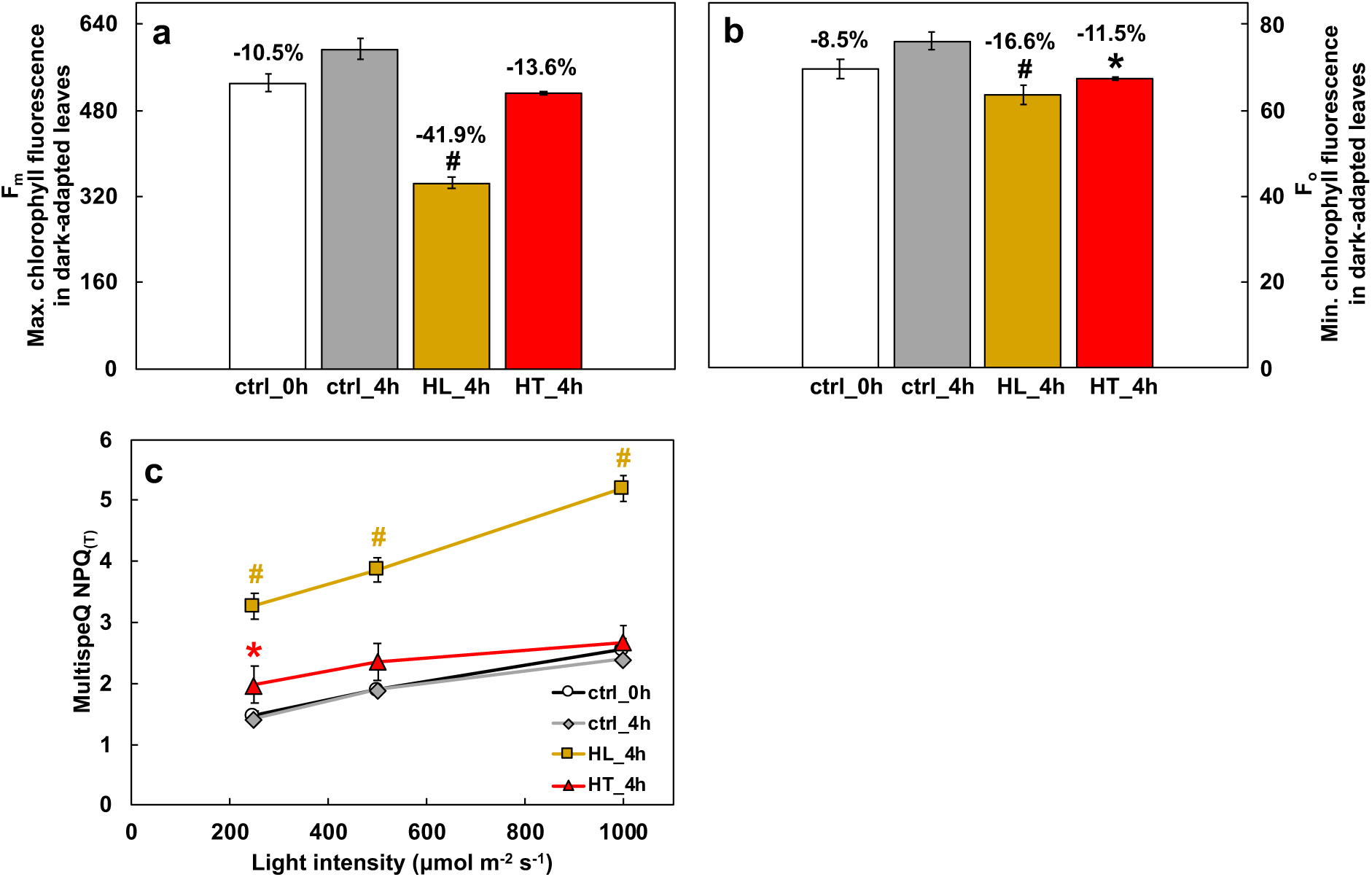
High light (HL) resulted in significant reduction of maximum chlorophyll fluorescence (F_m_) and the HL-induced NPQ was confirmed by using MultispeQ. **(a)** HL treatments resulted in significantly reduced maximal chlorophyll fluorescence in 20 min dark-adapted leaves (F_m_), however, F_m_ in ctrl_4h leaves were consistent among replicates. **(b)** HL and HT treatments resulted in reduced minimum chlorophyll fluorescence in dark-adapted leaves (F_o_) but F_o_ in ctrl_4h leaves were consistent among replicates. Percentages indicate reduction in F_m_ or F_o_ compared to ctrl_4h. **(c)** Estimated Non-photochemical quenching, NPQ_(T)_, calculated by F_o_’ and F_m_’ obtained during light response using MultispeQ. F_o_’ and F_m_’ are minimum and maximum chlorophyll fluorescence in light-adapted leaves. Mean ± SE, *n* = 3-6 biological replicates. Asterisk and pound symbols indicate statistically significant differences of ctrl_0h (at the start of treatments), HL_4h (after 4 h HL), and HT_4h (after 4 h HT) compared to ctrl_4h (after 4 h control treatment) using Student’s two-tailed t-test with unequal variance (*0.01<p<0.05, #p<0.01). For panel c, the colors of * and # match the significance of the indicated conditions, yellow for HL_4h, red for HT_4h).

**Supplementary Figure 10.**
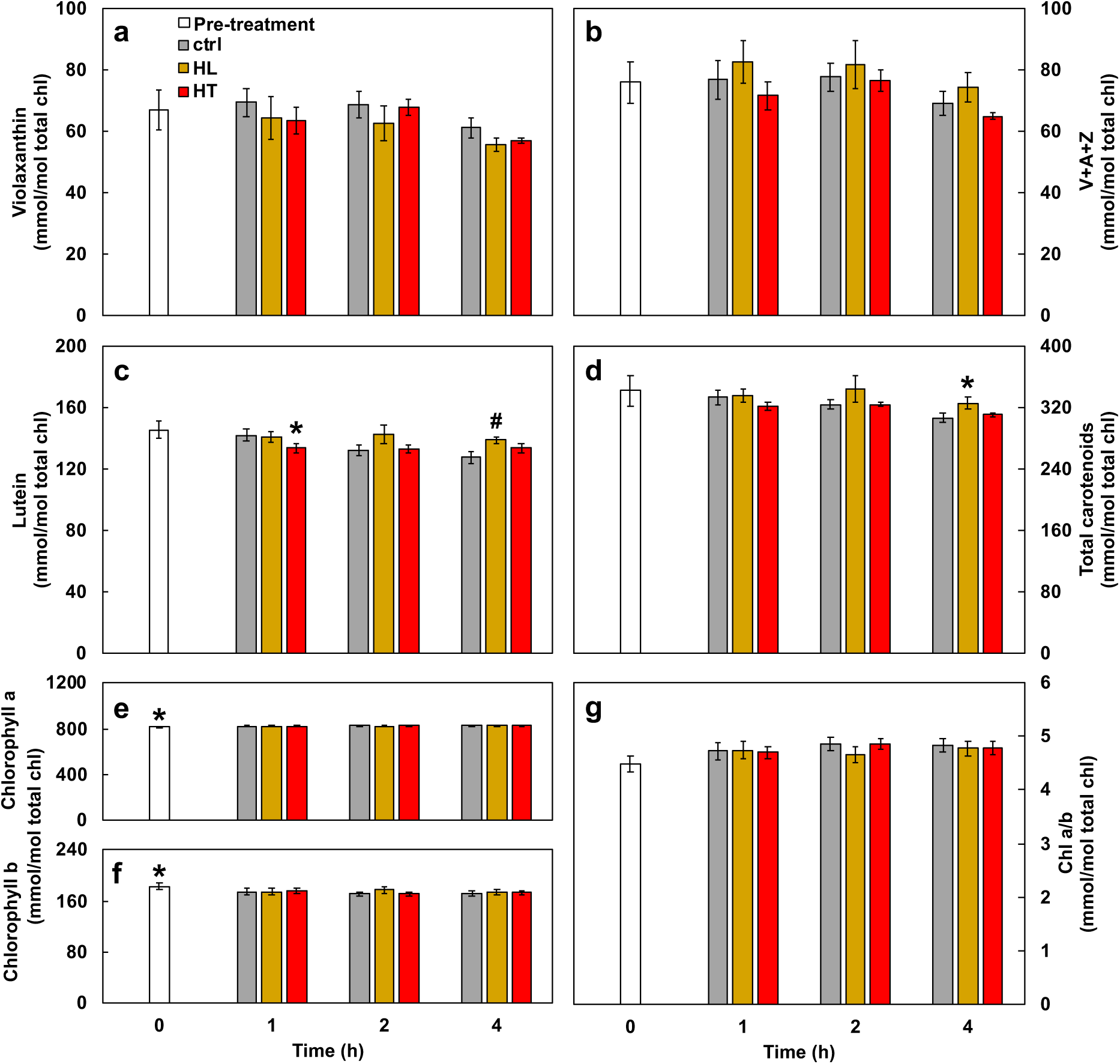
High light treatment increased lutein and carotenoids formation. Leaves of *S. viridis* were harvested for high-performance liquid chromatography (HPLC) analysis before treatment or after 1, 2, 4 h treatments of control growth condition (ctrl, 31°C and 250 μmol m^−2^ s^−1^), or high light (HL, 31°C and 900 μmol m^−2^ s^−1^) or high temperature (HT, 40°C and 250 μmol m^−2^ s^−1^). (a) Violaxanthin. (b) Total xanthophyll pool (violaxanthin + antheraxanthin + zeaxanthin, V+A+Z). (c) Lutein. (d) Total carotenoids. (e,f) Chlorophyll a and b. (g) Chlorophyll a/b ratio. Mean ± SE, *n* = 3 biological replicates. *0.01<p<0.05, #p<0.01, compared to control leaves at the same time points. Students’ two-tailed t-test with unequal variance.

**Supplementary Figure 11.**
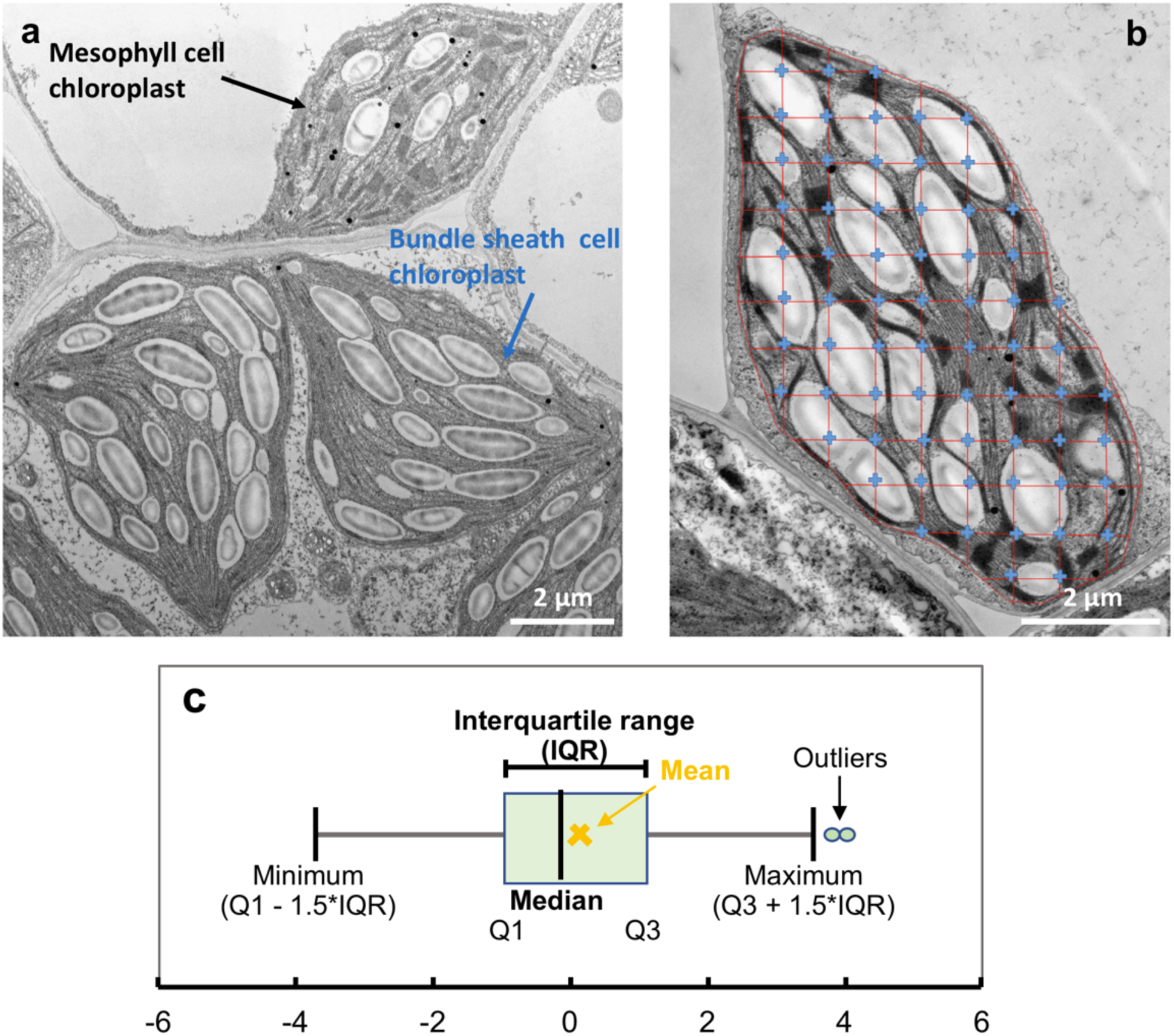
Representative transmission electron microscopy (TEM) images illustrate two cell types in *S. viridis* leaves, Stereo Analyzer analysis to quantify chloroplast structures, and boxplot of TEM data. **(a)** TEM image of chloroplasts of the two cell types in *S. viridis*: mesophyll cells and bundle sheath cells. **(b)** Illustration of Stereo Analyzer analysis for TEM images, which was used to calculate the relative volume of a cellular structures, e.g. starch granules. The Stereo Analyzer outlines a chloroplast with equally spaced uniform grid within the outlined area. The blue crossings of the grid inside the chloroplast are identified as either starch granule, stroma, stroma lamellae, or grana when they overlap with these structures. When all crossings have been identified, the software provides the % of relative volume for each structure of interest. **(c)** Illustration of TEM boxplots based on Tukey-style whiskers. Q1, first quartile; Q3, third quartile; IQR, interquartile range. The median value is represented by the vertical black line between Q1 and Q3. The mean value is represented by the yellow X sign.

**Supplementary Figure 12.**
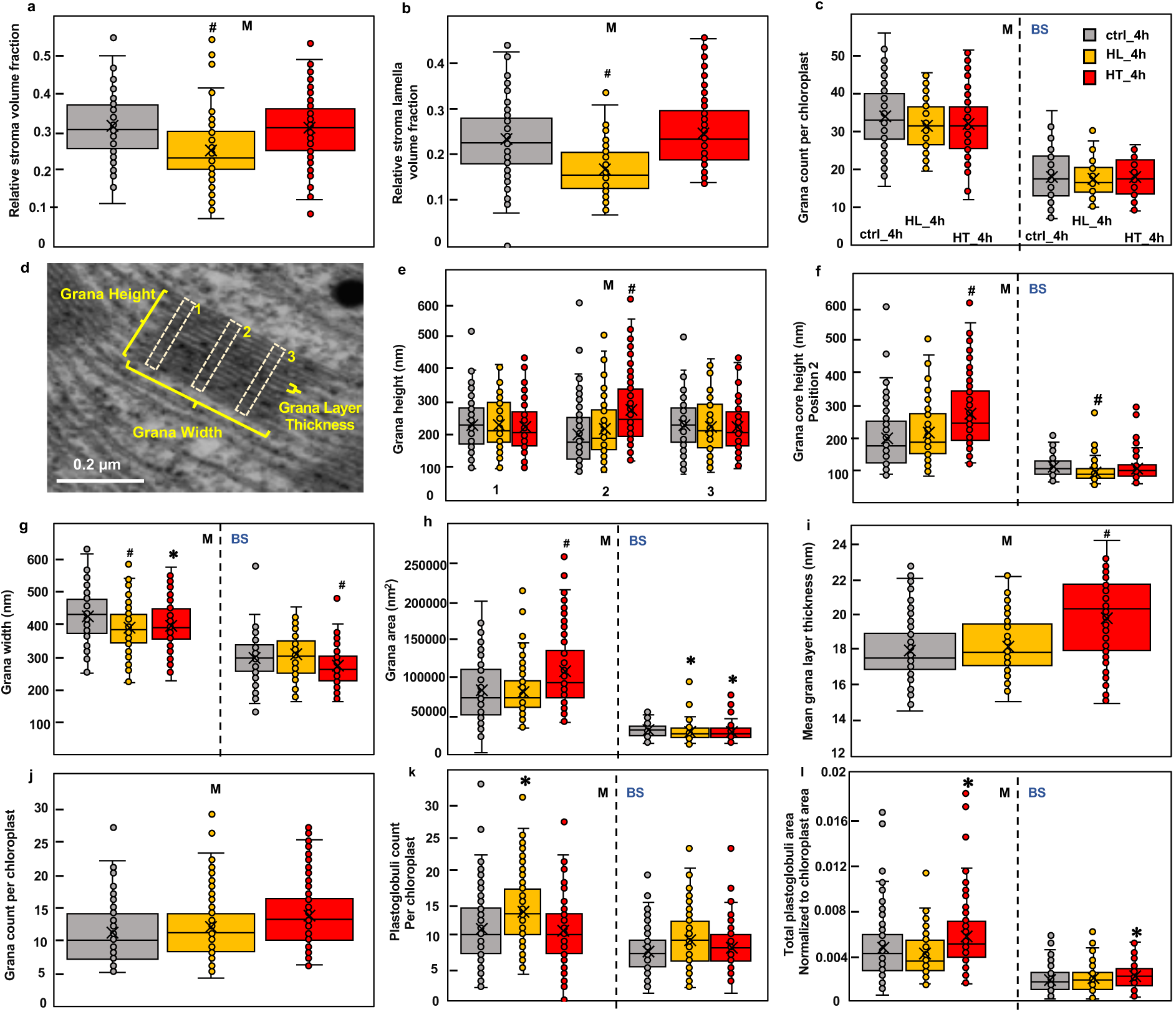
High light or high temperature altered various chloroplast structures in *S. viridis* leaves. Chloroplast structure changes after 4 h different treatments of control growth condition (ctrl_4h, 31°C and 250 μmol m^−2^ s^−1^), or high light (HL_4h, 31°C and 900 μmol m^−2^ s^−1^) or high temperature (HT_4h, 40°C and 250 μmol m^−2^ s^−1^). M, mesophyll chloroplast; BS, bundle sheath chloroplast. (a, b) Relative volume fractions were quantified using Stereo Analyzer with Kolmogorov–Smirnov test for statistical analysis compared to the same cell type of the control condition. (e, f, g, l) Parameters related to size and area were quantified using ImageJ with two-tailed t-test with unequal variance compared to the same cell type of the control condition. (c, j, k) The counting data was quantified using ImageJ followed by the negative binomial test for significance compared to the same cell type of the control condition. (d, e) Position 1 and 3 on grana are to measure the height of grana margin and position 2 is to measure the height of grana core. (h) Assuming grana are rectangular, grana area was estimated as grana core height multiplied by grana width. (i) The mean grana layer thickness was calculated as grana core height divided by the number of grana layers. Each treatment had three biological replicates with 90-120 images.

**Supplementary Figure 13.**
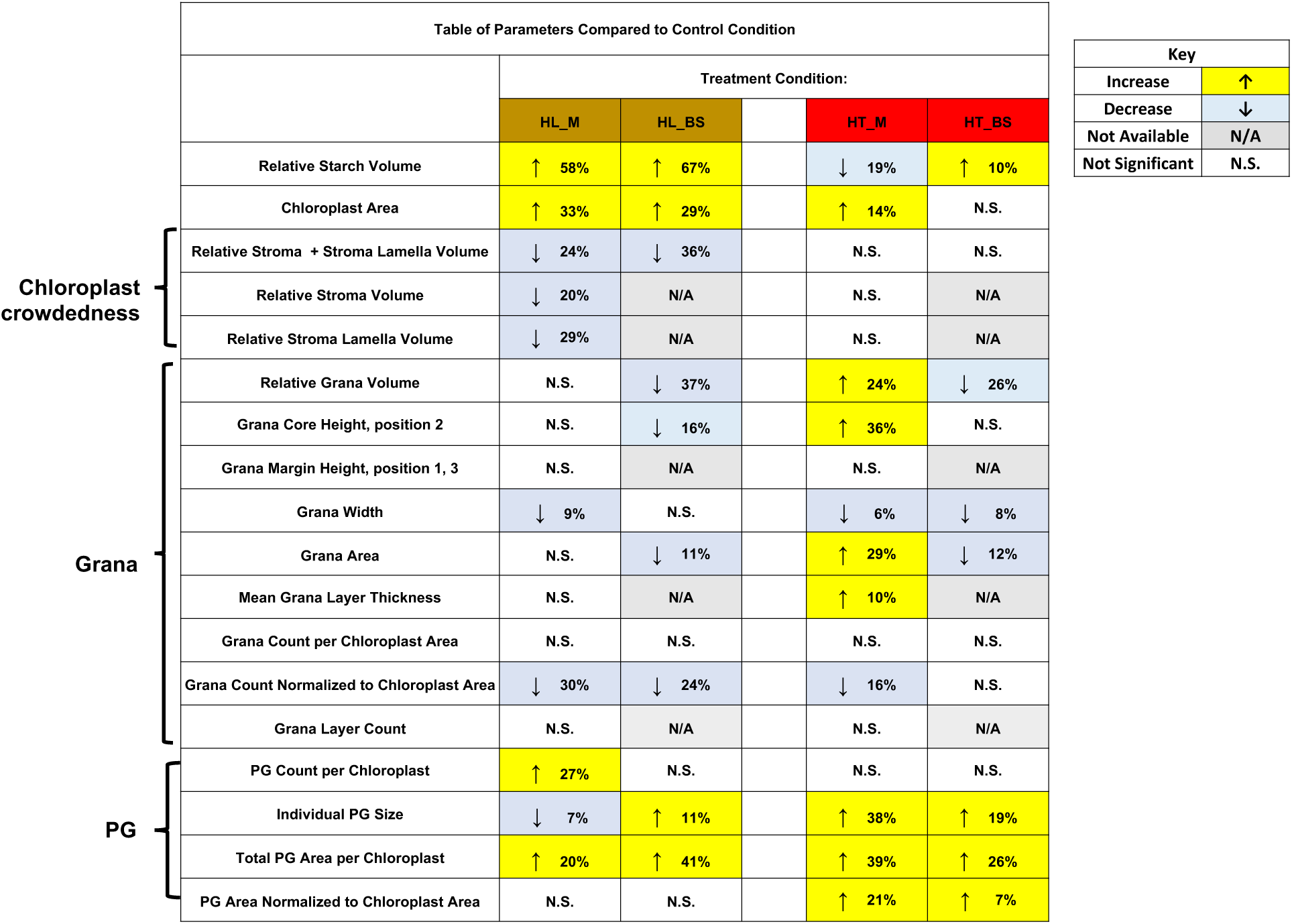
**Summary of chloroplast structure changes by using transmission electron microscopy (TEM) images** in leaves after 4 h treatments of high light (HL, 31°C and 900 μmol m^−2^ s^−1^) or high temperature (HT, 40°C and 250 μmol m^−2^ s^−1^) as compared to the control growth condition (ctrl, 31°C and 250 μmol m^−2^ s^−1^). BS, bundle sheath chloroplast; M, mesophyll chloroplast. PG, plastoglobuli. Mean value of each parameter was used for comparison. Yellow highlighted cells and upward arrows denote increased percentages as compared to the control condition. Blue highlighted cells and downward arrows denote decreased percentage as compared to the control condition. Grey highlighted cells and N/A mean data unavailable due to the difficulties to quantify some structures in the BS chloroplasts. White cells and N.S. mean no significant differences between HL or HT as compared to the control treatment.

**Supplementary Figure 14.**
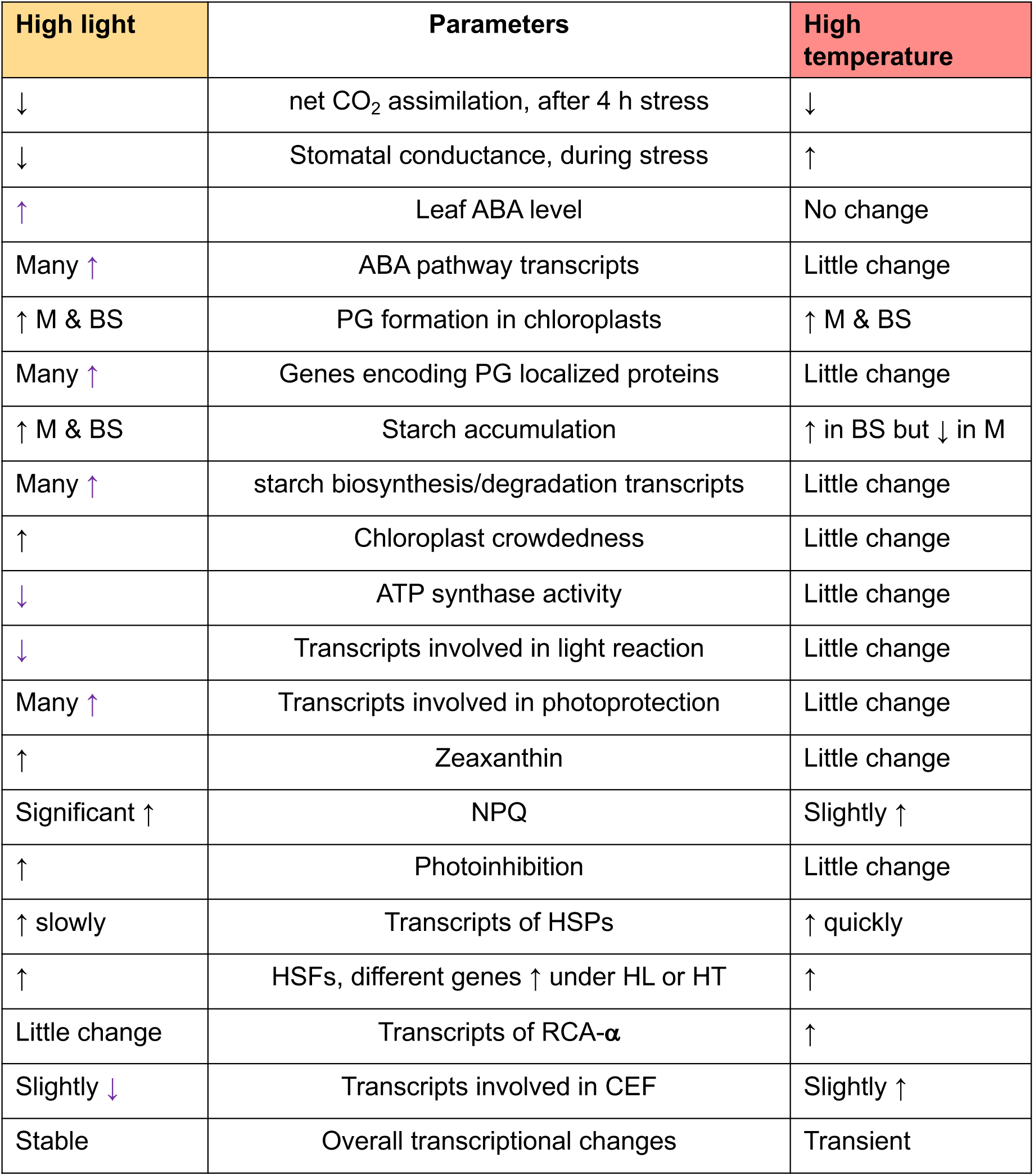
Summarized multi-level changes of *S. viridis* in response to 4 h high light or high temperature treatments as compared to the control treatment. Upward arrows denote increase or induction; downward arrows denote decrease or repression. HL, high light; HT, high temperature; ABA, abscisic acid; M, mesophyll chloroplast; BS, bundle sheath chloroplast; PG, plastoglobuli; NPQ, non-photochemical quenching; HSP, heat shock protein; HSF, heat shock transcription factor; RCA, Rubisco activase; CEF, cyclic electron flow around PSI.

