## Supplementary_Table_1_photosynthetic_parameters for "High Light and High Temperature Reduce Photosynthesis via Different Mechanisms in the C_4_ Model *Setaria viridis*"

| Description | Label | Units | Formula |
|---|---|---|---|
| **Net $CO_2$ assimilation rate** | $A_{Net}$ | µmol m$^{-2}$ s$^{-1}$ | $A = \frac{Flow(CO_{2_R} - CO_{2_S}\left(\frac{1000 - H_2O_R}{1000 - H_2O_S}\right))}{100S}$ <br> $Flow$: air flow rate (µmol s$^{-1}$) <br> $CO_{2_R}$: reference cell $CO_2$ concentration (µmol mol$^{-1}$) <br> $CO_{2_S}$: sample cell $CO_2$ concentration (µmol mol$^{-1}$) <br> $H_2O_R$: reference cell $H_2O$ mole fraction (mmol mol$^{-1}$) <br> $H_2O_S$: sample cell $H_2O$ mole fraction (mmol mol$^{-1}$) <br> $S$: leaf area in cm$^2$ |
| **Transpiration rate** | $E$ | mol m$^{-2}$ s$^{-1}$ | $E = \frac{Flow(H_2O_S - H_2O_S)}{1000S(1000 - H_2O_S)}$ |
| **Stomatal conductance to water vapor** | $g_{sw}$ | mol m$^{-2}$ s$^{-1}$ | $g_{sw} = \frac{2}{\left(\frac{1}{g_{tw}} - \frac{1}{g_{bw}}\right) + \sqrt{(\frac{1}{g_{tw}} - \frac{1}{g_{bw}})^2 + \frac{4K}{(K+1)^2}(2\frac{1}{g_{tw}} - \frac{1}{g_{bw}})\frac{1}{g_{bw}}}}$ <br> $g_{tw}$: total conductance of the leaf to water vapor (mol m$^{-2}$ s$^{-1}$) <br> $g_{bw}$: boundary layer conductance to water vapor (mol m$^{-2}$ s$^{-1}$) <br> $K$: stomatal ratio |
| **Intercellular $CO_2$** | $C_i$ | µmol mol$^{-1}$ | $C_i = \frac{\left(g_{tc} - \frac{E}{2}\right) CO_{2_S} - A}{g_{tc} + \frac{E}{2}}$ |
| **Maximal chlorophyll fluorescence, dark-adapted leaves** | $F_m$ | | |
| **Maximal chlorophyll fluorescence, light-adapted leaves** | $F_m'$ | | |
| **Minimal chlorophyll fluorescence, dark-adapted leaves** | $F_o$ | | |
| **Minimal chlorophyll fluorescence, light-adapted leaves** | $F_o'$ | | |
| **Steady state fluorescence** | $F_s$ | | |
| **Variable chlorophyll fluorescence** | $F_v$ | | $F_V = F_m - F_o$ |
| **PSII maximum efficiency in dark-adapted leaves** | $F_v/F_m$ | | ${F_v}/{F_m} = {F_v}/{F_m} = 1 - \frac{F_o}{F_m}$ |
| **Non-photochemical quenching** | NPQ | | $NPQ = \frac{(F_m - F_m')}{F_m'}$ |
| **Estimated NPQ** | $NPQ_{(T)}$ | | $NPQ_{(T)} = \left(\frac{4.88}{\frac{F_m'}{F_o'} - 1}\right) - 1$ |
| **PSII operating efficiency** | ΦPSII | | $\Phi PSII = 1 - \frac{F_s}{F_m'}$ |
| **Electron transport rate** | ETR | µmol m$^{-2}$ s$^{-1}$ | $ETR = (\Phi PSII)(0.5)(Qabs_{fs})$ <br> $Qabs_{fs}$: absorbed light corresponding to the last $F_s$ measurement |
| **Fraction of open PSII centers** | $q_L$ | | $q_L = q_P * \frac{F_o'}{F_s}$ |
| **Plastoquinone redox status** | $Q_A$ | | $Q_A = 1 - q_L$ |
