## Supplementary material for "High Light and High Temperature Reduce Photosynthesis via Different Mechanisms in the C_4_ Model *Setaria viridis*": SupFig11_with_high_resolution_TEM_image

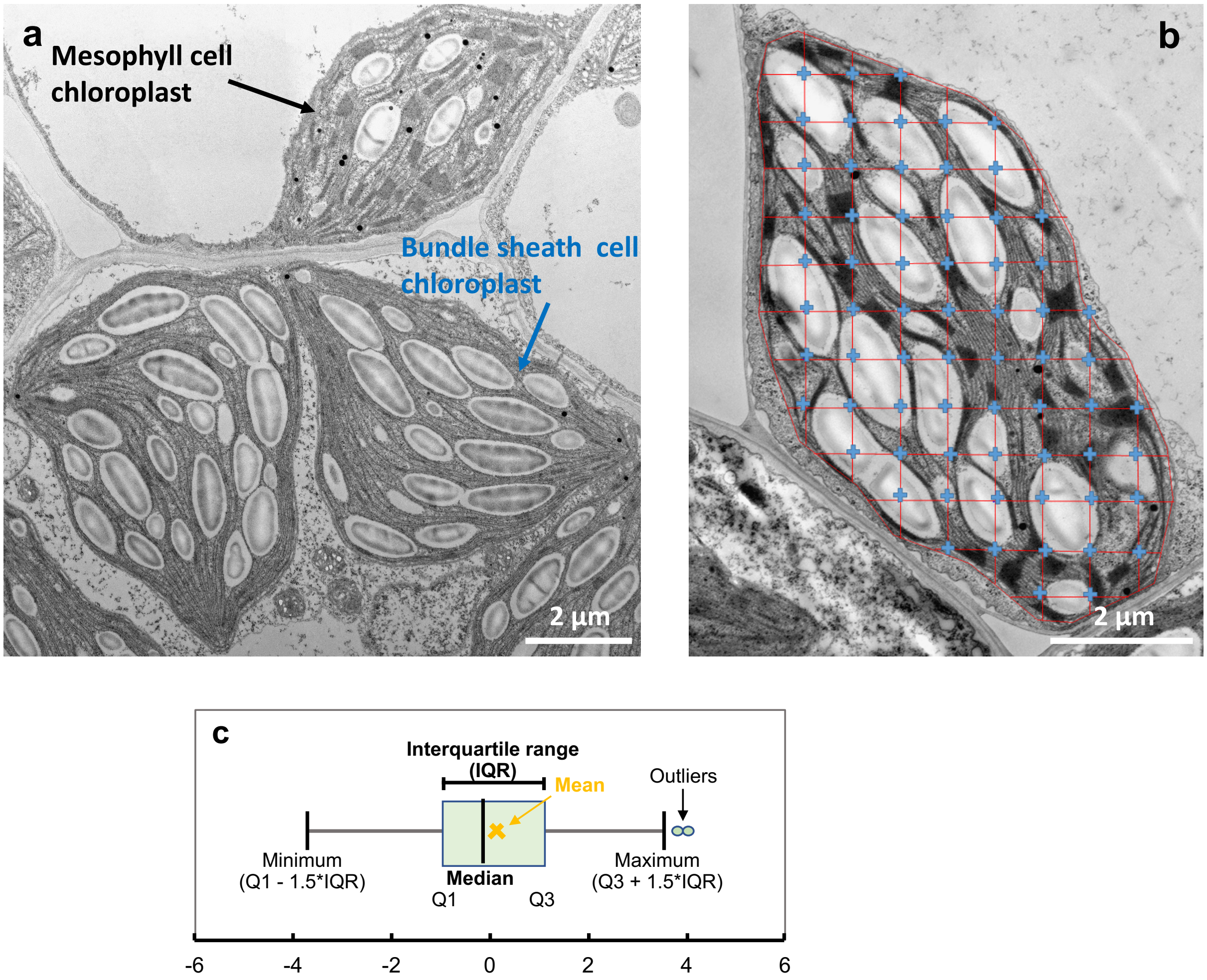


**Supplementary Figure 11. Representative transmission electron microscopy (TEM) images illustrate two cell types in *S. viridis* leaves, Stereo Analyzer analysis to quantify chloroplast structures, and boxplot of TEM data. (a)** TEM image of chloroplasts of the two cell types in *S. viridis*: mesophyll cells and bundle sheath cells. **(b)** Illustration of Stereo Analyzer analysis for TEM images, which was used to calculate the relative volume of a cellular structures, e.g. starch granules. The Stereo Analyzer outlines a chloroplast with equally spaced uniform grid within the outlined area. The blue crossings of the grid inside the chloroplast are identified as either starch granule, stroma, stroma lamellae, or grana when they overlap with these structures. When all crossings have been identified, the software provides the % of relative volume for each structure of interest. **(c)** Illustration of TEM boxplots based on Tukey-style whiskers. Q1, first quartile; Q3, third quartile; IQR, interquartile range. The median value is represented by the vertical black line between Q1 and Q3. The mean value is represented by the yellow X sign.
