## Supplementary material for "High Light and High Temperature Reduce Photosynthesis via Different Mechanisms in the C_4_ Model *Setaria viridis*": Fig7_with_high_resolution_TEM_images

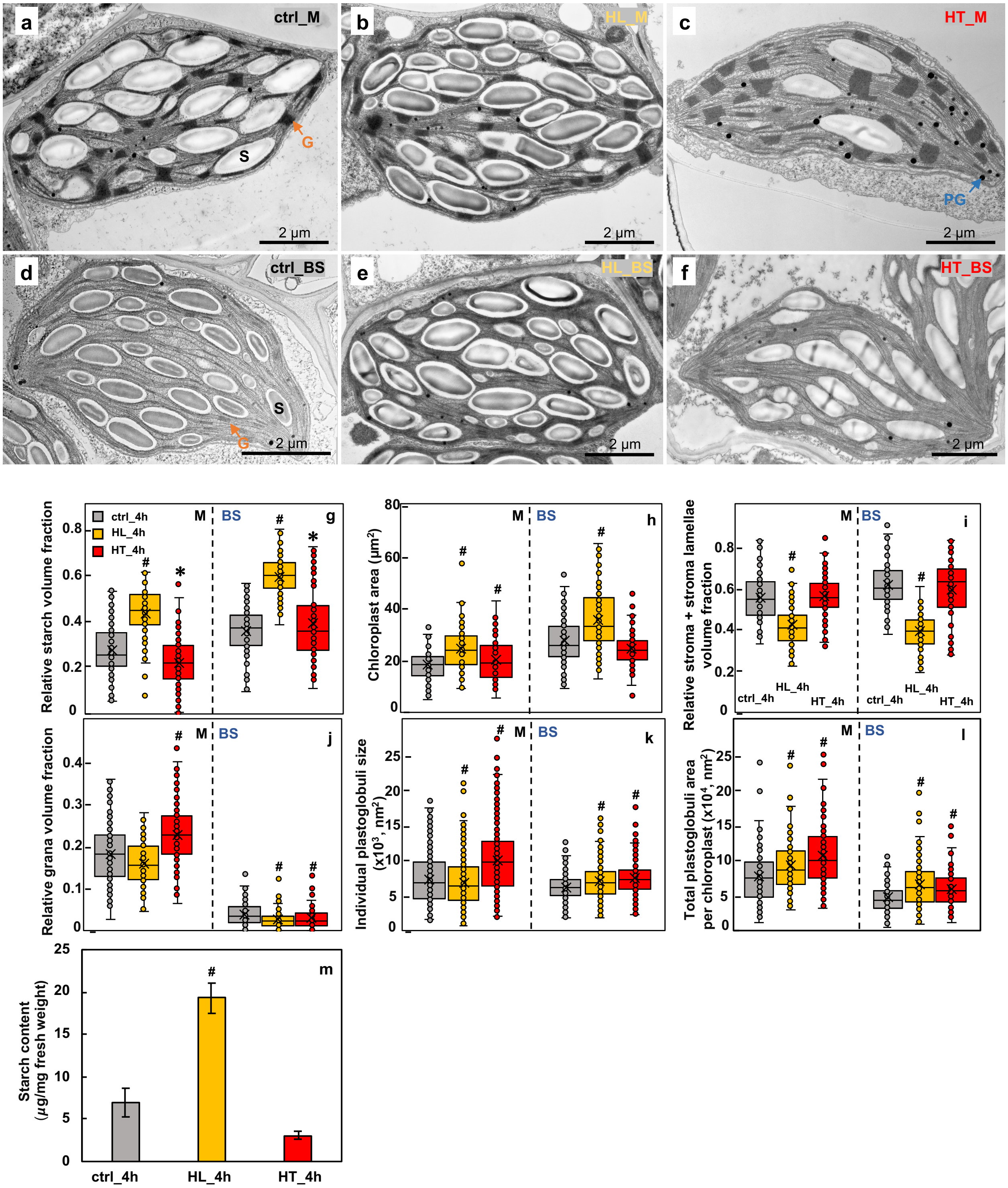


**Figure 7: High light (HL) increased starch accumulation and both HL and high temperature (HT) treatments induced chloroplast plastoglobuli formation in *S. viridis* leaves. (a-f)** Representative transmission electron microscopy (TEM) images of mesophyll (M) and bundle sheath (BS) chloroplasts in leaves of *S. viridis* after 4 h

treatments of control (ctrl_4h, 31°C, 250 μmol $m^{-2}$ $s^{-1}$ light) or high light (HL_4h, 31°C, 900 μmol $m^{-2}$ $s^{-1}$) or high temperature (HT_4h, 40°C, 250 μmol $m^{-2}$ $s^{-1}$ light). TEM images of M **(a, b, c)** and BS **(d,e,f)** chloroplasts. S labels the starch granule; G labels grana, the orange arrows indicate grana in M and BS chloroplasts; PG labels plastoglobuli. **(g, i, j)** Relative volume fraction of indicated parameters were quantified using Stereo Analyzer with Kolmogorov–Smirnov test for statistical analysis compared to the same cell type of the control condition. **(h, k, l)** area and size of indicated parameters were quantified using ImageJ with two-tailed t-test with unequal variance compared to the same cell type of the control condition. Each treatment had three biological replicates, total 90-120 images per treatment. *0.05<p<0.01; #p<0.01. (**m**) Starch quantification using starch assay kits. HL_4h leaves accumulated 4x starch as compared to ctrl_4h leaves. Values are mean ± SE, *n* = 3 biological replicates. Pound symbols indicate statistically significant differences as compared to ctrl_4h using Student's two-tailed t-test with unequal variance (#p< 0.01).
